# Human milk variation is shaped by maternal genetics and impacts the infant gut microbiome

**DOI:** 10.1101/2023.01.24.525211

**Authors:** Kelsey E. Johnson, Timothy Heisel, Mattea Allert, Annalee Fürst, Nikhila Yerabandi, Dan Knights, Katherine M. Jacobs, Eric F. Lock, Lars Bode, David A. Fields, Michael C. Rudolph, Cheryl A. Gale, Frank W. Albert, Ellen W. Demerath, Ran Blekhman

## Abstract

Human milk is a complex mix of nutritional and bioactive components that provide complete nutrition for the infant. However, we lack a systematic knowledge of the factors shaping milk composition and how milk variation influences infant health. Here, we used multi-omic profiling to characterize interactions between maternal genetics, milk gene expression, milk composition, and the infant fecal microbiome in 242 exclusively breastfeeding mother-infant pairs. We identified 487 genetic loci associated with milk gene expression unique to the lactating mammary gland, including loci that impacted breast cancer risk and human milk oligosaccharide concentration. Integrative analyses uncovered connections between milk gene expression and infant gut microbiome, including an association between the expression of inflammation-related genes with IL-6 concentration in milk and the abundance of *Bifidobacteria* in the infant gut. Our results show how an improved understanding of the genetics and genomics of human milk connects lactation biology with maternal and infant health.

## Introduction

Lactation is the defining trait of mammals and has been essential for our species for most of human evolution^1^. Today, breastfeeding is recommended as the exclusive mode of feeding for infants, given its documented health benefits for both mothers and infants^2^. The nutritional significance of human milk stems from hundreds of milk constituents, including macro- and micro-nutrients, immune factors, hormones, oligosaccharides, and microbes^3^. Maternal factors such as diet, health status, and genetics shape variation in milk composition across lactating women^4^; however, the relative importance of these factors on most milk components are poorly understood^5^. The role of maternal genetics in shaping milk composition is particularly understudied. A small number of studies suggest important relationships between maternal genotype, milk composition, and infant health^6^. For example, maternal secretor status, determined by the *FUT2* gene, is linked to human milk oligosaccharide (HMO) composition^7^. HMOs are sugars in human milk that cannot be digested by the infant but promote the growth of beneficial microbes in the infant gut, and may provide additional immunological and metabolic benefits^8^. In addition to HMOs, variation in other milk components, such as fatty acids, has been linked to the infant gut microbiome^9,10^, and breastfeeding (vs. formula feeding) is one of the strongest factors shaping the infant gut microbiome^11,12^. The abundance of certain microbes in the infant gut, particularly *Bifidobacteria*, has been linked to health outcomes in infancy and later childhood^13^. Thus, the composition of the infant gut microbiome represents a key outcome through which human milk promotes infant health. Here, we combine maternal clinical and milk composition data with maternal whole-genome sequences, milk transcriptomes, and infant fecal metagenomics to characterize genetic influences on gene regulation in milk and identify pathways linking milk gene expression with milk composition and infant gut health. The results advance our knowledge of the complex molecular and physiological relationships connecting mother, milk, and infant^14^.

### Milk gene expression correlates with maternal traits and milk composition in a healthy, successfully lactating cohort

Human milk contains mammary epithelial luminal cells and a variety of immune cell types, including macrophages, lymphocytes, and granulocytes^15–19^. Thus, a milk sample provides rich information on the biology of milk production and immune phenotypes in the lactating mammary gland^15,16^. To characterize population-level variation in human milk gene expression, we performed bulk RNA-sequencing on the cell pellets from 1-month postpartum milk samples from 242 women in the Mothers and Infants LinKed for Healthy Growth (MILK) study^20–22^ (Fig. S1-3, Table S1). Comparison to gene expression data from human tissues obtained by the GTEx consortium^23^ showed that milk expression profiles clustered near other secretory tissues, such as the pancreas, kidney, and colon (**Fig. 1A**, Fig. S4). The three most highly expressed milk genes (*CSN2, LALBA, CSN3*), which comprise a large proportion of milk transcripts^15^, accounted for 34.5% of protein-coding transcripts in milk, reminiscent of the preponderance of hemoglobin transcripts typical in whole blood (**Fig. 1B**)^23^. These three genes encode the major milk proteins beta- and kappa-casein (*CSN2, CSN3*) and lactalbumin (*LALBA*), an essential protein for lactose and HMO synthesis^24^.

**Figure 1.**
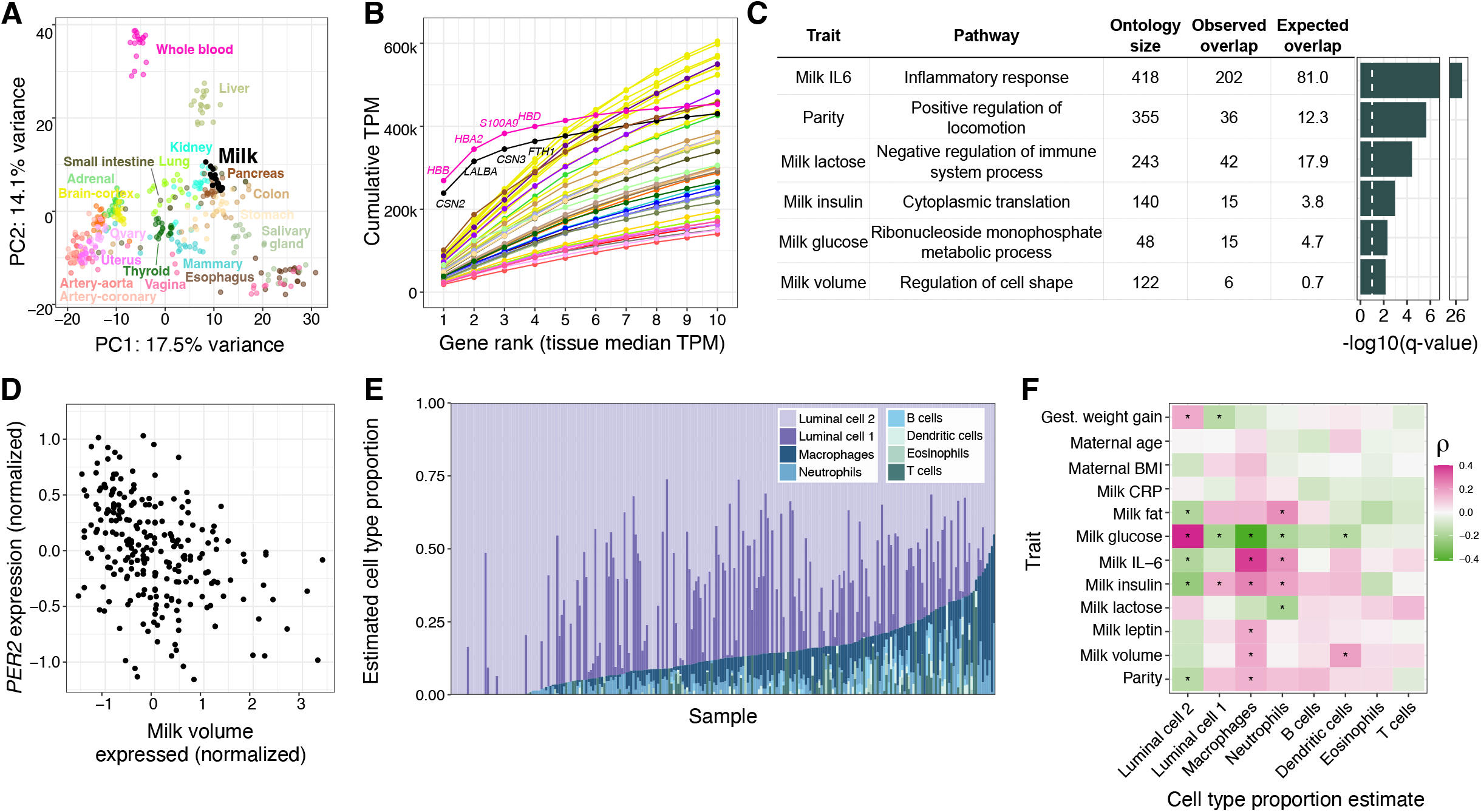
Overview of gene expression in human milk. **A)** Principal components analysis of transcriptomes from a subset of GTEx tissues and milk. PCs were calculated using the 1000 most variable genes within GTEx, then milk samples were projected onto the GTEx samples. An equivalent plot including all GTEx tissues is in Fig. S1. **B)** Cumulative TPM (transcripts per million) of the top 10 genes by median TPM for milk and GTEx tissues. Color scheme is the same as in 1A. **C)** Gene ontology enrichment of genes with expression correlated to maternal and milk traits. The most significant term for each trait is shown (Methods). The dashed white vertical line denotes a q-value of 10%. **D)** Correlation between milk volume (from a standardized electric breast pump expression during a study visit, see Methods) and normalized *PER2* gene expression in milk. **E)** Cell type proportion estimates generated using Bisque^30^ for transcriptomes from this study, and reference milk single cell RNA-seq from Nyquist et al 2022^17^. **F)** Heatmap of Spearman correlations between estimated cell type proportions (x-axis) and maternal/milk traits (y-axis). *q-value<10%.

To identify factors associated with the milk transcriptome, we tested for correlations between the expression of 12,584 genes in milk and 12 maternal or milk traits (Table S2-3, Fig. S5). Among maternal traits, only parity (the number of previous births) was significantly correlated with expression of at least one gene (423 genes at q-value<10%; negative binomial generalized log-linear test, see Methods). Genes for which expression correlated with parity were enriched for pathways related to cell locomotion, potentially reflecting persistent differences in mammary gland remodeling during lactation in participants who had previously lactated^25^ (**Fig. 1C**). Pre-pregnancy BMI and gestational weight gain, traits associated with delayed lactogenesis and breastfeeding challenges^26^, were not significantly correlated with milk gene expression (Table S3). This lack of relationship could be due to our study’s inclusion of only women who successfully breastfed for at least 1 month postpartum, thus excluding participants with difficulties initiating breastfeeding related to metabolic health. Milk concentrations of IL-6, glucose, insulin, and lactose were each correlated with expression of hundreds of genes, and the total single breast milk expression volume produced at the study visit was correlated with 65 genes (q-value<10%; Table S3). These milk trait-correlated genes were enriched for processes such as cytoplasmic translation (milk insulin) and regulation of cell shape (milk volume) (**Fig. 1C**, Table S4).

The gene for which expression was most significantly associated with expressed milk volume is the core circadian clock gene *PER2*. Higher *PER2* expression correlated with lower milk volume (**Fig. 1D**), and was also correlated with a higher percentage of milk fat (Table S3). The relationship between *PER2* expression and milk volume or milk fat was not simply driven by the time of day of milk expression (volume: ANOVA P=0.77; fat: P=0.75). In addition to *PER2*, the expression levels of 4 of 21 genes in the circadian rhythm pathway were nominally associated (P<0.05) with milk volume (*PER1, PER3, NPAS2, FBXL3;* Table S3). *PER2* plays a role in cell fate and ductal branching in the mammary gland^27^, and clock gene expression rhythms are suppressed in the mammary gland during lactation, possibly to enable milk production in response to suckling cues^28^. Our observation suggests that differential expression of circadian clock genes in the mammary gland affects milk production in humans, possibly via regulation of milk production genes or by anatomical changes in the breast during lactogenesis.

Of all milk traits tested, IL-6 protein concentration was correlated with expression of the largest number of genes (2,291 genes at q-value<10%; Table S3). Genes positively correlated with milk IL-6 concentration were enriched for immune pathways, with “inflammatory response” the most significantly enriched pathway (q-value = 2.9×10^-27^, Fisher’s exact test; **Fig. 1C**), consistent with IL-6’s role as a marker of inflammation in the mammary gland^29^. To estimate the contributions of different cell types to our milk bulk transcriptomes, we performed cell-type deconvolution using a milk single cell RNA-seq reference panel (**Fig. 1E**; Methods)^17,30^. Consistent with previous studies, mammary epithelial cells were estimated to make up the majority of cells^17–19,31^. The estimated proportion of neutrophils and macrophages were increased in milk samples with higher IL-6 concentration (neutrophils: multiple regression coefficient = 0.32, q-value = 8.4×10^-4^; macrophages: multiple regression coefficient = 0.24, q-value =1.5×10^-3^; **Fig. 1F;** Table S5), suggesting the relationship between IL-6 concentration and immune gene expression is caused by a greater proportion of immune cells in milk.

### Genetic influences on gene expression in human milk

Associations between genetic variation and gene expression can illuminate the molecular mechanisms underlying genetic influences on human traits^32^, but this approach has not been applied to human milk. To identify associations between maternal genetic variation and milk gene expression, we generated low-pass whole genome sequencing data and performed an expression quantitative trait locus (eQTL) scan in 206 unrelated human milk samples (Methods). We identified a local eQTL (q-value<5%) at 2,690 genes out of 16,999 tested (Table S6). Comparing milk eQTLs to those identified in 45 human tissues in the GTEx project^23^, we partitioned our eQTLs as milk-specific (N=487) or shared with at least one other tissue (N=2,203) (**Fig. 2A**; Table S6). Genes with milk-specific eQTLs highlighted key biological pathways in the lactating mammary gland: production of caseins (e.g. the abundant milk proteins *CSN3 and CSN1S1);* lactose synthesis (*LALBA*); lipogenesis (e.g. *ACSL1, CD36, LPL, LPIN1, SCD5, SPTLC3);* hormonal regulation (*INSR);* and immunity (e.g. *LYZ, MUC7, CD68*) (Table S6). In addition, genes with milk-specific eQTLs were twice as likely as genes with eQTLs shared across multiple tissues to overlap genetic associations for milk traits in dairy cattle (odds ratio = 2.1, P-value = 1.7×10^-3^, Fisher’s exact test; **Fig. 2B**; Table S7), a species for which there is far more known about genetic influences on lactation than in humans. This enrichment suggests that genes with milk-specific eQTLs are specifically important for milk biology. Genes with milk-specific eQTLs also tended to have more sequence-level constraint^33^ than tissue-shared eQTLs (P-value = 1.3×10^-7^, Wilcoxon rank sum test; **Fig. 2C**), and were enriched for the pathways “regulation of ERK1 and ERK2 cascade” and “long-chain fatty-acyl-CoA metabolic process” (**Fig. 2D**, Methods). These pathways are physiologically relevant in milk, as ERK cascade signaling has a key role in mammary morphogenesis^34^, and lipogenesis generates the energy dense fats synthesized by the lactating mammary gland^35^.

**Figure 2:**
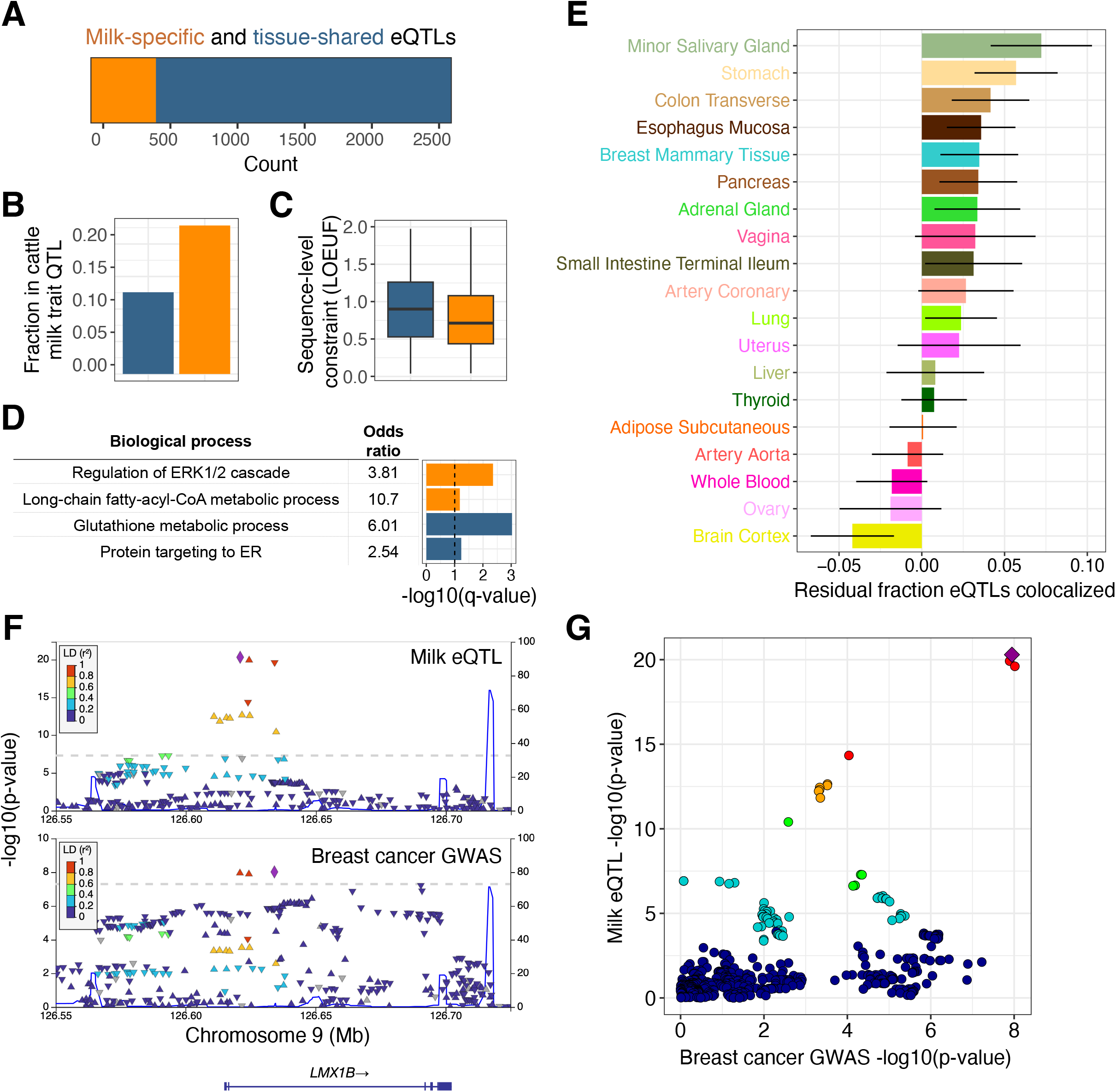
Genetic influences on gene expression in human milk. **A)** Counts of genes that have milk-specific eQTLs (orange, genes that have an eQTL only in milk or where the milk eQTL did not colocalize with any GTEx tissue, see Methods) vs. tissue-shared eQTLs (blue, genes with milk eQTLs that colocalized with at least one other tissue in GTEx). **B)** Fraction of genes in each category that overlapped with a milk trait QTL in the dairy cattle genome. **C)** Distributions of sequence-level constraint, measured by the loss-of-function observed/expected upper bound fraction (LOEUF) statistic^33^. **D)** Enriched gene ontologies for genes with milk-specific (orange) or tissue-shared (blue) eQTLs. The dashed vertical line denotes a q-value of 10%. **E)** Sharing of eQTLs between milk and a subset of GTEx tissues, measured through statistical colocalization. Each bar shows each tissue’s similarity to milk, measured by the residual fraction of eQTLs colocalized with milk, after regressing out tissue sample size. Error bars represent a 95% confidence interval. **F)** LocusZoom genetic associations in the *LMX1B* region with milk gene expression (top panel) and breast cancer risk (bottom panel). Each data point represents a SNP, plotted by their chromosomal location (x-axis) and significance of association (y-axis), with colors corresponding to LD (linkage disequilibrium, r^2^) to the lead SNP for each dataset, shown as a purple diamond. **G)** Each point is a variant, plotted by the strength of association with milk gene expression (y-axis) and breast cancer risk (x-axis). Colors are the same as the top panel in 2F, with a purple diamond representing the lead milk eQTL SNP. The pattern of variants in the top right suggests a shared underlying causal variant.

To identify tissues for which genetic regulation of gene expression is most similar to milk, we measured the proportion of shared eQTLs between milk and each GTEx tissue. After correction for tissue sample size, milk shared the largest proportion of eQTLs with secretory tissues (minor salivary gland, stomach, and colon), with a higher proportion shared than that observed for non-lactating breast tissue (**Fig. 2E**, Fig. S6). These comparisons highlight the shared regulation of gene expression across secretory epithelial tissues, and underscore the insufficiency of resting breast tissue for studying gene expression programs necessary for lactation.

Epidemiological studies describe a complex relationship between lactation and breast cancer risk, with increased short-term risk associated with pregnancy, but decreased lifetime risk associated with longer duration of lactation^36^. Because the genetics of gene expression in the lactating mammary gland is distinct from that of resting breast (**Fig. 2E**), milk eQTLs provide unique functional annotations to genetic associations with breast cancer. Using colocalization analyses between all milk eQTLs and breast cancer GWAS loci, we identified 9 loci with strong evidence for a shared causal variant (posterior probability of shared causal variant > 0.9; Table S8). Of these milk eQTL-GWAS colocalizations, 8 had previously been nominated as a causal gene for breast cancer^37–40^. We identified a novel candidate gene for one breast cancer GWAS locus, where a milk-specific eQTL that increased expression of *LMX1B* was associated with increased cancer risk (**Fig. 2F, 2G**). *LMX1B* is a transcription factor essential for normal development of limbs, kidneys, and ears^41^.

### Milk gene expression correlates with concentrations of human milk oligosaccharides

Maternal genetics play a strong role in shaping the concentration of HMOs^7^, sugars in milk that are not digested by the infant but promote the growth of beneficial microbes in the infant gut. HMOs are synthesized in the mammary gland by addition of monosaccharides to a lactose molecule, but the glycosyltransferases catalyzing these reactions are largely uncharacterized^42^. Secretor status, determined by the absence of a common nonsense variant in the fucosyltransferase 2 (*FUT2*) gene, strongly predicts the concentration of certain HMOs, with the presence of some HMOs entirely determined by secretor status^7^. Utilizing 48 participants with both milk gene expression and 1-month HMO composition data, we observed distinct HMO profiles between secretors and non-secretors (**Fig. 3A**, Fig. S7; see Table S9 for HMO definitions). We hypothesized that beyond the strong effects of the secretor polymorphism, the expression of *FUT2* in milk would correlate with HMO concentrations within secretor individuals, reflecting variation in milk among women with a functional FUT2 enzyme. We observed nominally significant associations between *FUT2* expression and the concentration of two HMOs: 2’FL (Beta = 0.40, P = 0.03; Fig. S8) and 6’SL (Beta = −0.42, P = 0.04; Fig. S8). This suggested that milk gene expression data could be useful for identifying critical genes for HMO biosynthesis. We tested for pairwise correlations between gene expression and 19 individual HMOs (Fig. S9), and the sums of all HMO concentrations, sialylated HMOs, and fucosylated HMOs. Of these 22 HMO traits, 14 were significantly correlated with expression of between 1 and 196 genes (q-value<10%, Table S10). These included known HMO biosynthesis genes, such as the sialyltransferase *ST6GAL1*^42^ with total HMO concentration (Beta = 0.75, P = 7.2×10^-5^, q-value=0.08; **Fig. 3B**). All HMO traits significantly correlated with the expression of more than ten genes were sialylated HMOs (Table S10). The genes correlated with sialylated HMO concentrations were enriched for inflammatory immune pathways, such as “cellular response to lipopolysaccharide” enriched in genes correlated with total sialylated HMO concentration (**Fig. 3C**, Table S11), consistent with previous evidence that the sialylated HMOs 6’SL, LSTc, and DSLNT were more abundant in women with mastitis compared to healthy women^43^.

**Figure 3.**
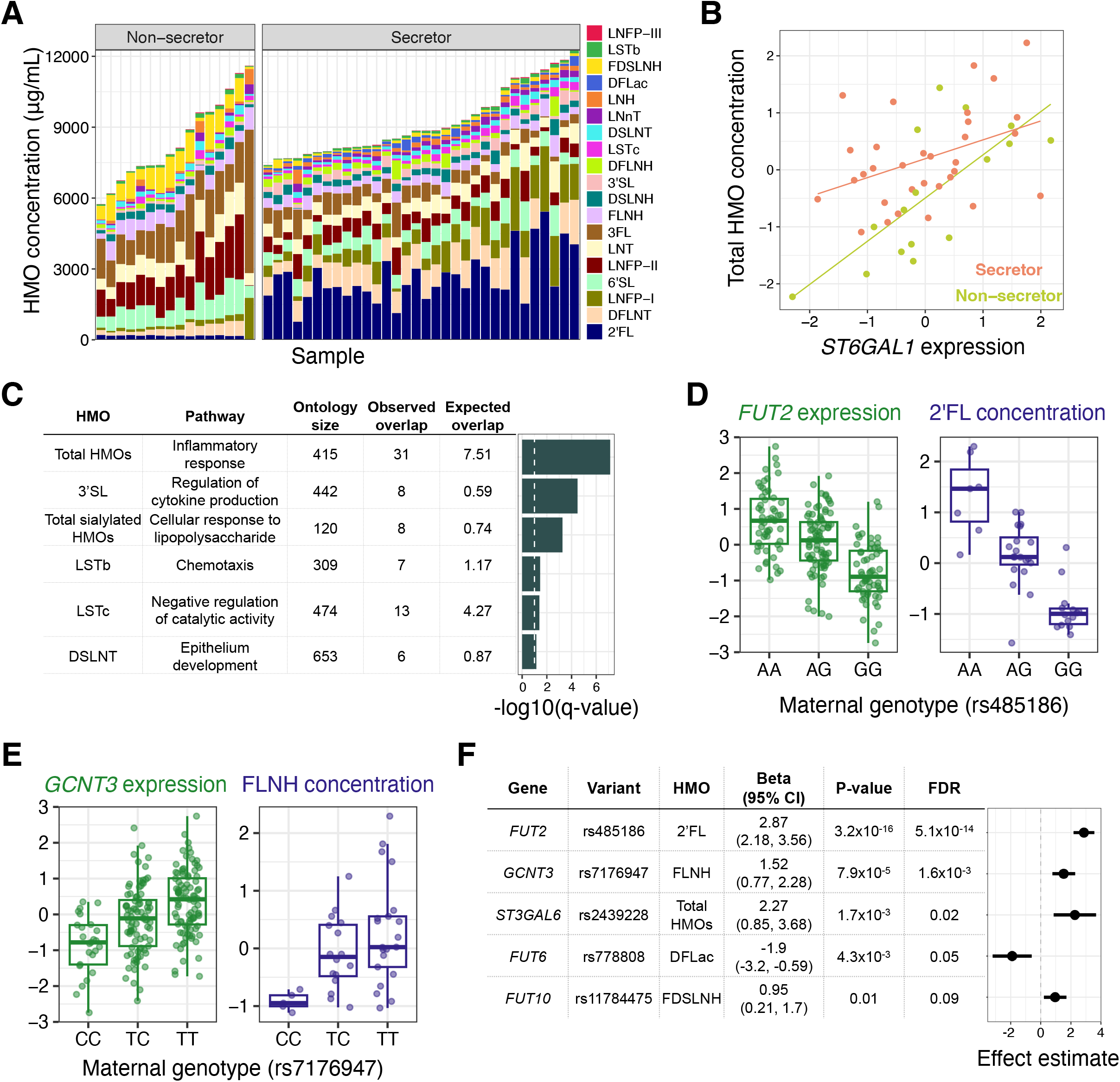
Effects of milk gene expression on HMO composition. **A)** HMO concentration profiles (y-axis) for milk samples in our study (x-axis), grouped by secretor status. **B)** Correlation between *ST6GAL1* gene expression in milk and normalized total HMO concentration, colored by secretor status (beta = 0.75, P = 7.2×10^-5^, q-value = 0.08.). **C)** Gene ontology enrichment of genes with expression correlated to a single HMO or HMO category. The most significant term for each HMO is plotted. The dashed vertical line denotes a q-value of 10%. **D)** Relationships between genotype at the lead SNP at the *FUT2* eQTL and *FUT2* expression in milk (green) or 2’FL abundance (purple). **E)** Relationships between genotype at the lead SNP at the *GCNT3* eQTL and *GCNT3* expression in milk (green) or FLNH abundance (purple). **F)** Estimates of the effect of milk gene expression of candidate HMO-biosynthesis pathway genes on the abundance of HMOs, from a Wald ratio test. Some genes had significant effects on more than one HMO (Table S11). The most significant HMO for each gene is plotted here.

HMO biosynthesis represents an ideal system to understand the effects of maternal genetics on milk composition via changes in gene expression, as gene expression from the relevant cell type (mammary epithelial cells) and HMO concentrations can be measured non-invasively in the same milk samples. Among 54 candidate glycosyltransferase genes^42^, eight genes had significant milk eQTLs in our data (Table S12), which we used to test for associations between maternal genotypes at milk eQTL tag SNPs and HMO concentrations. For five genes we observed an association between genotype and between 1 and 12 HMOs (Table S13; q-value<10%). These included the known association of *FUT2* with 2’FL (**Fig. 3D**), and an association between *GCNT3* and FLNH (**Fig. 3E**). *GCTN3* was also linked to FLNH in our above analysis of correlations between gene expression and HMO concentrations (Table S10, Fig. S10). *GCTN3* was identified previously as the best candidate gene responsible for the addition of a β-1,6-linked N-acetylglucosamine to the lactose core, a step required for the biosynthesis of FLNH^42^. For each of 160 eQTL-HMO pairs, we then estimated the causal effect of modified gene expression on HMO concentration using a Wald ratio test, and found a significant effect in 18 eQTL-HMO pairs (**Fig. 3F**; q-value<10%, Table S13). These results provide evidence for direct or indirect roles of specific glycosyltransferases in HMO biosynthesis in the lactating mammary gland.

### Maternal genotype and milk gene expression is associated with the infant gut microbiome

Studies have found correlations between milk composition and variation in the infant gut microbiome^9,10,44^. However, it is unclear how these correlations are shaped by maternal genetics and milk gene regulation. We hypothesized that given milk gene expression reflects milk composition, it could be correlated with the infant gut microbiome. We profiled the fecal microbiome of infants in our study with metagenomic sequencing at 1 (N=108) and 6 (N=113) months postpartum (**Fig. 4A**, Fig. S11), and identified six correlated sets of genes expressed in milk and microbial taxa or pathways present in the infant gut at 1 month postpartum using sparse canonical correlation analysis^45^ (sparse CCA, see Methods; **Fig. 4B**, Table S14). Using pathway enrichment analysis, we identified relevant biological processes in these milk-expressed gene sets correlated with the infant fecal microbiome. For example, milk expression of T-cell receptor signaling genes was negatively correlated with the abundance of *Haemophilus spp*. in the infant gut (**Fig. 4C**), and expression of N-glycan biosynthesis pathway genes in milk was negatively correlated with bacterial ketogluconate metabolism pathway abundances (**Fig. 4D**). These links between milk gene expression and the infant gut microbiome nominate biological pathways through which normal, healthy variation in human milk composition influences the infant gut microbiome.

**Figure 4.**
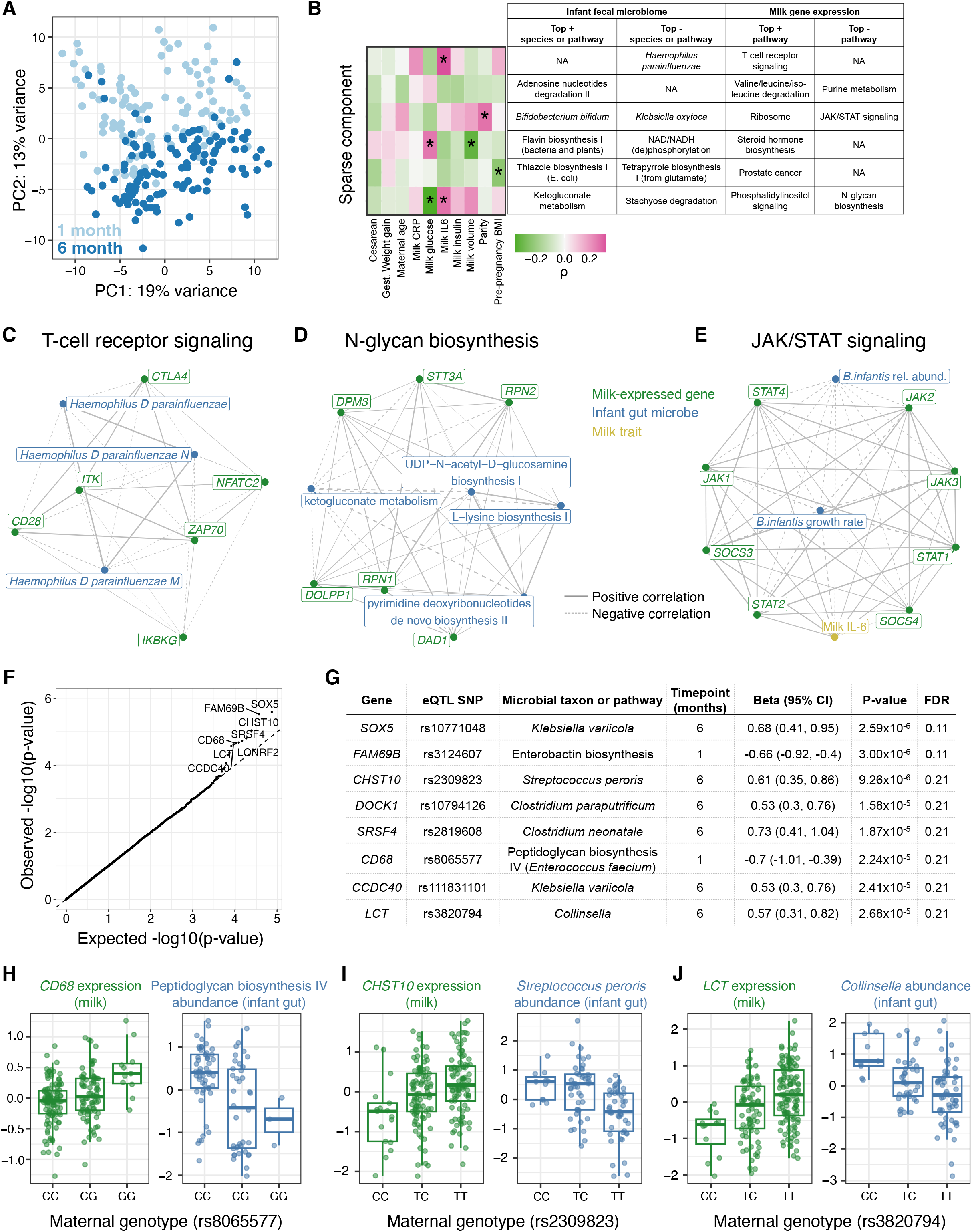
Interactions between milk gene expression and the infant fecal microbiome. **A)** Principal components analysis of infant fecal microbiome metagenomic data, summarized at the taxonomic level, with each point representing a fecal sample and colors representing infant age (light blue: 1 month; dark blue: 6 months). **B)** Sparse canonical correlation analysis integrating milk host gene expression and infant fecal microbial species or microbial gene pathway relative abundances (at 1 month of age) identified six significant sparse components (in rows). The heatmap on the left shows correlation coefficients between each mother/infant pairs’ score for a given sparse component and clinical data (in columns). The table lists the top most highly weighted microbial taxon or genetic pathway, and most significantly enriched host gene set in milk gene expression. (+) or (-) indicates if these features were positively or negatively weighted in the sparse component. **C-D)** Network diagrams generated using the correlation matrix of infant fecal microbial species/pathways and milk-expressed host genes within an enriched pathway for two of the sparse components in (B). Line size corresponds to the absolute value of correlation coefficient, line type correspond to negative (dashed) or positive (solid) correlations. Node color signifies milk-expressed host genes (green), infant fecal microbial pathways/taxa (green), or milk traits (yellow). **E)** Network diagram displaying correlations between milk IL-6 concentration, JAK/STAT pathway genes expressed in milk, and *Bifidobacterium infantis* relative abundance and estimated growth rate in the infant gut at 1 month. JAK/STAT pathway genes were selected that had a significant correlation with either *B. infantis* trait after multiple test correction (q-value<10%). **F)** Q-Q plot showing expected (x-axis) vs. observed (y-axis) p-values from association tests between maternal genotype at milk-specific eQTLs and relative abundances of infant fecal microbial taxa/pathways. Top associations are labeled with the gene name. **G)** Details on 8 associations (rows) between milk eQTL and infant fecal microbe abundance that passed q-value<25%. **H-I)** Associations between maternal genotype at a milk-specific eQTL with the expression of that gene in milk (green, left), and with the relative abundance of an infant microbiome feature (blue, right).

The sparse CCA algorithm identified species of *Bifidobacterium* in the infant gut as correlated with milk-expressed genes in the JAK/STAT pathway, which is a key regulator of both milk production and mammary inflammation^46^. Given our observation that genes in this pathway were significantly correlated with milk IL-6 concentration (Table S3), we further examined the relationships between milk expression of JAK/STAT pathway genes, milk IL-6 concentration, and infant fecal *Bifidobacterium infantis*, including computationally-inferred *B. infantis* growth rates (Methods). *B. infantis* is an abundant microbe in the breastfed infant gut that promotes beneficial health outcomes^47,48^. Both infant fecal *B. infantis* growth rate and relative abundance were negatively correlated with milk expression of JAK/STAT pathway genes, most significantly *STAT1* (growth rate: Pearson’s r=-0.70, P=9.7×10^-5^; relative abundance: r=-0.24, P=0.02; **Fig. 4E**, Table S15). *STAT1* encodes a key element of the mammary anti-inflammatory response to bacterial mastitis^49^ and is mainly expressed in the immune cells in milk^17^. Thus, the correlation between increased *STAT1* signaling in milk and lower *B. infantis* abundance and growth in the infant gut could be related to an immune response to infection of the mammary gland.

Finally, we tested for associations between maternal genotypes at milk-specific eQTLs and infant gut microbiome traits (**Fig. 4F**, Table S16), reasoning that such associations could be mediated through differences in milk composition. While no associations were significant at the q-value<10% level, we identified 8 potential associations between maternal genotype and infant fecal microbiome with q-value<25% (**Fig. 4G**). These included a milk-specific eQTL for the macrophage marker gene *CD68*, at which the expression-increasing allele was associated with lower abundance in the 1 -month infant gut of the microbial pathway “peptidoglycan biosynthesis IV” in species of *Enterococci* (**Fig. 4H**). At an eQTL for *CHST10*, the expression-increasing allele was associated with lower *Streptococcus peroris* abundance in the 6-month infant gut (**Fig. 4I**). The enzyme encoded by *CHST10* (HNK-1 sulfotransferase) participates in the synthesis of glycosaminoglycans (GAGs)^50^. GAGs are abundant in human milk^51^ and prevent pathogenic bacterial adhesion to epithelial cells^52,53^; and lower infant gut *Streptococcus peroris* is associated with decreased diarrhea risk^54^. We also found an association between a milk-specific eQTL at the lactase (*LCT*) gene with infant gut genus *Collinsella* at 6 months (**Fig. 4J**). The milk *LCT* expression-increasing allele, which also increases lactase expression in the intestines of European adults^55^, is correlated with decreased infant gut *Collinsella*. This eQTL was detected as ‘milk-specific’ in our study because *LCT* had no significant eQTL in any GTEx tissue (q-value<5%). Maternal *LCT* genotype could alter the breastfed infant microbiome via changes in milk composition, maternal diet, and/or the maternal microbiome.

## Discussion

Here, we generated and integrated multiple omics datasets within a cohort of exclusively breastfeeding mother-infant pairs, leveraging the milk transcriptome as a readout of the biology of milk production. Our results highlight how an improved understanding of the genetics and genomics of human milk reveals connections with maternal and infant health.

A consistent theme across our results was a link between mammary inflammation-related gene expression, milk composition, and the infant gut microbiome. Milk IL-6 concentration explained the most variation in milk gene expression of all tested traits (Table S3). Genes correlated with the concentration of sialylated HMOs in milk were enriched for inflammation-related pathways (**Fig. 3C**, Table S11); and expression of JAK/STAT pathway genes in milk, particularly *STAT1*, were inversely correlated with the abundance and growth of *Bifidobacteria* in the infant gut (**Fig. 4E**). All participants in our study were exclusively breastfeeding and did not report symptoms of mastitis (infection of the mammary gland) at the time of milk collection. Thus, our results suggest that mammary inflammation, even when unnoticeable to the lactating individual, is a primary driver of variation in milk composition, with potential effects on the infant gut microbiome.

Combining milk gene expression with maternal genetic variation, we identified numerous novel milk-specific eQTLs, which can now be used as targets for investigation of the effects of gene expression on milk production and composition, and infant and maternal health. For example, combining our milk eQTLs with breast cancer GWAS summary statistics, we provide the first functional evidence connecting *LMX1B* expression to a nearby breast cancer GWAS locus (**Fig. 3F,3G**). Functional evidence for this GWAS locus had previously been missing, as this milk-specific eQTL may only be detectable during lactation. In an analysis of single cell RNA-seq across human tissues, *LMX1B* was most highly expressed in salivary and breast glandular cells^56^. In addition, hypomethylation at a CpG island in *LMX1B* in human milk samples was associated with subsequent diagnosis of breast cancer in an epigenome-wide association study^57^, suggesting higher expression correlated with breast cancer risk, which is concordant with the direction of effect in our results.

We also show that milk eQTLs can be leveraged to understand the effects of milk gene expression on the breastfed infant. We identified an intriguing association between an allele near the *LCT* gene that confers higher lactase expression in milk and decreased infant gut *Collinsella* (**Fig. 4G,4J**). The same allele confers lactase persistence in adults, a phenotype that is likely to have provided selective advantage in periods of famine and/or infectious disease during human evolution^58^. The lactase persistence allele is replicably associated with differences in the adult gut microbiome^59^, but has not previously been linked to the microbiome of infants. Moreover, a recent paper found evidence of an adaptive advantage of the lactase-persistence allele in British infants during WWII through an analysis of infant mortality^60^. Our results raise the possibility that maternal genetic effects on infants, possibly mediated by milk composition, could be under selection at this locus. The *LCT* eQTL does not have an effect until later in childhood post-weaning^61^, such that an indirect (maternal) genetic effect rather than a direct effect of infant genotype provides a plausible mechanism for selection on breastfed infants who have not yet experienced the age-dependent effects of the lactase persistence allele. *LCT* is expressed in mammary cells at levels comparable to intestine^56^, though no role for the lactase enzyme in milk production has been described. More work is needed to understand how the *LCT* genotype impacts milk composition, and its effects on the infant gut microbiome. Looking forward, large cohorts with both maternal and infant genetic information and rich phenotyping will be needed to assess potential effects of maternal genotype on infant health mediated by milk composition.

While our study introduced a framework for integrating multiple and diverse data types in the mother/milk/infant triad, it is limited by the sample sizes of our milk composition phenotypes (especially HMOs) and infant fecal microbiome data. We are also hindered by the lack of infant genotypes in our study, which may account for some of the observed maternal genetic associations with the infant gut microbiome. Additionally, the MILK study is predominantly composed (~85%) of participants who self-identify as white and non-Hispanic. Thus, our analysis was limited to genetic variants common in participants of European ancestry, and our eQTL results may not be generalizable to other ancestral groups. Lastly, we studied mature milk collected at 1-month postpartum which did not allow us to assess genetic effects on colostrum or milk produced at other points in lactation.

The importance of breastfeeding, especially in underdeveloped countries, is widely acknowledged, but the long-term health effects in modern high-income contexts are less concrete^2^. Similarly, the causal effects of differences in milk composition for breastfed infants are underexplored due to the ethical and logistical impediments to performing randomized trials of infant nutrition. The field of human genetics has been hugely successful in identifying genetic effects on molecular and complex traits, and has leveraged these associations to improve understanding of disease pathophysiology, identify drug candidates, and interrogate causal relationships impacting human health. However, traits related to women’s health generally have been overlooked by this area of research, and human milk and lactation is a glaring example of this neglect. Fortunately, milk represents an easily obtained non-invasive biospecimen, aiding our ability to close this gap. Our study provides a step towards leveraging modern human genomics techniques to characterize the factors that shape milk composition, understand how this composition impacts infant and maternal health, and eventually utilize the information to support policy and behavioral interventions to optimize breastfeeding and breastmilk at the population level.

## Acknowledgements

We thank Katy Duncan, Laurie Foster, Tipper Gallagher, and all MILK study staff and participants for their contributions, and members of the Albert and Blekhman labs for helpful discussions related to this project. This work was supported by the resources and staff at the University of Minnesota Genomics Center (https://genomics.umn.edu). This work was carried out in part by resources provided by the Minnesota Supercomputing Institute (https://www.msi.umn.edu/).

## Author contributions

Conceptualization: KEJ, FWA, EWD, RB

Formal analysis: KEJ, TH, MA

Funding acquisition: KEJ, DK, KMJ, EFL, LB, DAF, CAG, FWA, EWD, RB

Investigation: KEJ, TH, AF, NY

Supervision: KEJ, LB, MCR, CAG, FWA, EWD, RB

Writing - original draft: KEJ

Writing - review and editing: KEJ, TH, EFL, LB, MCR, CAG, FWA, EWD, RB

## Data and materials availability

Non-identifiable data (RNA-seq quantifications, infant fecal metagenomic abundances, HMO concentrations, milk eQTL summary statistics, relevant metadata) is available at figshare (https://figshare.com/projects/Johnson_et_al_human_milk_multi-omics/156606). Genotype and milk RNA sequencing data will be deposited at dbGaP, and metagenomic sequencing data at the SRA, prior to publication.

## List of supplementary materials

Materials and Methods

Figs. S1 to S11

Tables S1 to S16

References (62–92)

