## Supplemental material for "Human milk variation is shaped by maternal genetics and impacts the infant gut microbiome"

K.E. Johnson, et al.
2023

### Materials and Methods

#### MILK Study overview

Participant recruitment, clinical data, and milk sample collection for the Mothers and Infants LinKed for health (MILK) study have been described previously[20–22,62,63]. Briefly, participants who intended to exclusively breastfeed were enrolled prenatally during healthy, uncomplicated pregnancies at the University of Minnesota in collaboration with HealthPartners Institute (Minneapolis, MN) or the University of Oklahoma Health Sciences Center. Recruited mothers were 21–45 years old, non-smokers, non-diabetic, and delivered singleton infants at full term (37 0/7 – 41 6/7 weeks gestation) with 10th–90th percentile birth weight on the WHO growth chart. No participants reported symptoms of mastitis or breast infection at the time of milk sample collection. Clinical data for each mother-infant dyad was collected from the delivering hospitals' electronic health record and from electronic questionnaires at study visits at 1 and 6 months postpartum. The Institutional Review Boards of the University of Oklahoma, the University of Minnesota, and the HealthPartners Institute approved this study (STUDY00009021). This study has been registered with ClinicalTrials.gov (identifier NCT03301753). The data described in this manuscript comes from a subset of MILK Study mother/infant pairs who consented to maternal whole-genome sequencing, milk RNA sequencing, and microbiome assessment of infant fecal samples.

Milk samples were collected at study visits at approximately 1 month postpartum, and infant fecal samples were collected at study visits at 1 and 6 months. Upon study visit arrival, participants fed their infants *ad libitum* from one or both breasts until infants were satisfied. Two hours following this feeding, milk was collected from the right breast using a hospital-grade electric breast pump (Medela Symphony; Medela, Inc., Zug, Switzerland), with expression ceasing when milk stopped flowing. Expressed milk volume and weight was recorded, milk was gently mixed, aliquots were made, and then stored at -80°C within 20 minutes of collection and kept at -80°C until thawed for RNA/DNA extraction.

#### Milk composition measurements

**Human milk oligosaccharides (HMOs)**: Concentrations of HMOs were quantified from 2 mL previously frozen whole milk aliquots as previously described[64]. 19 HMOs were identified and quantified: 2′-fucosyllactose (2'FL), 3-fucosyllactose (3'FL), 3′-sialyllactose (3'SL), 6′-sialyllactose (6'SL), difucosyllactose (DFLac), difucosyllacto-N-hexaose (DFLNH), difucosyllacto-N-tetrose (DFLNT), disialyllacto-N-hexaose (DSLNH), disialyllacto-N-tetraose (DSLNT), fucodisialyllacto-N-hexaose (FDSLNH), fucosyllacto-N-hexaose (FLNH), lacto-N-fucopentaose (LNFP) I, LNFP II, LNFP III, lacto-N-hexaose (LNH), lacto-N-neotetraose (LNnT), lacto-N-tetrose (LNT), sialyl-lacto-N-tetraose b (LSTb), and sialyl-lacto-N-tetraose c (LSTc). Secretor milk was defined as having a 2′FL concentration that was greater than a natural, very low break in the data (Fig. 3A). Weight-based concentrations were used for all statistical analyses (micrograms per milliliter). The sum of HMO concentrations was calculated as the total concentrations of the 19 measured HMOs.

**Milk cytokines/nutrients/hormones**: Milk fat was separated from the aqueous phase by centrifugation, and skim milk was assayed using commercially available immunoassay kits for insulin, glucose, leptin, CRP, and IL6 as previously described[20,22,65]. These milk component assays were processed in 2-5 batches depending on the assay. Batch effects were corrected using an analysis of variance model with formula

log(assay value)~factor(batch)

using the 'aov' command in R. The residuals from this model, representing the batch-corrected values, were used in all downstream data analyses.

**Milk fat and lactose**: Milk fat and lactose concentrations were assessed using mid-infrared spectrophotometry (Calais Milk Analyzer, North American Instruments, LLC, Lake Oswego, OR)[66,67]. Human milk samples were gradually thawed and then diluted with deionized water in a 1:1 dilution. Breastmilk control samples with standard macronutrient content were run prior to study sample testing to confirm instrument calibration. Samples were heated in a water bath until the samples reached 40°C and were mixed by gentle hand inversion for 2 minutes prior to analysis, per manufacturer instructions. Milk fat percent reliability was assessed in a random subset of 34 samples (17 duplicate samples) with an intraclass correlation coefficient (ICC) of 0.99, $p<0.001$. Validity was assessed in a random subset of 30 samples against the gold standard Mojonnier method[68] yielding a high cross-method ICC of 0.936, $p<0.001$.

### RNA extraction and sequencing

We extracted RNA from whole milk cell pellets to capture gene expression from both mammary epithelial cells and immune cells in milk. Previous studies that have performed bulk RNA-sequencing from human milk have used RNA extracted from the milk fat layer[15]. This procedure enriches for milk fat globule RNA, which originates from mammary epithelial cells[15,16]. Our approach allowed us to use computational approaches to estimate the contribution of different cell types to our milk transcriptome, and explore genetic influences on gene expression that could be specific to the immune cells in milk, in addition to mammary epithelial cells.

Nucleic acid extractions and RNA-seq library preparation and sequencing was performed at the University of Minnesota Genomics Center (UMGC). Frozen 2 mL whole milk aliquots from 245 milk samples were thawed and split in two, with each 1mL half used for either RNA or DNA extraction. RNA was extracted from the cell pellet using the RNeasy Plus Universal HTP following the manufacturer's instructions. We used the TakaraBio Stranded Total RNA Pico Mammalian kit for RNA-seq library preparation. RNA libraries were sequenced on an Illumina NovaSeq 6000 S2 flow cell with 2x150 paired-end reads to a median depth of 34.2 million reads per sample (Table S1).

### RNA-seq pre-processing and quantification

RNA-seq reads were trimmed with Trimmomatic and aligned with STAR[69] v2.7.1a to the GRCh38 human reference genome. Gene-level quantification was performed with RNA-SeQC[70] v2.3.4 using a Gencode v36 gene model annotation that was collapsed to a single transcript model per gene using a script provided by GTEx ("collapse_annotation.py" from https://github.com/broadinstitute/gtex-pipeline/tree/master/gene_model).

To assess the gene-level quantifications, TPM spearman correlations were calculated between each pair of samples with the ‘rcorr’ function from the ‘Hmisc’ R package[71]. One outlier sample was removed for having low median pairwise correlations with other samples and fewer than 10,000 genes detected (Fig. S1). There were two participants that each contributed two milk samples, from two separate pregnancies. We included only one milk sample from each of these participants in our analyses, leaving 242 milk transcriptomes from 242 different participants (Table S1).

#### Whole-genome sequencing and quality control

DNA was extracted from the cell pellet using the QIAamp 96 DNA Blood Kit at UMGC following the manufacturer’s instructions. Low-pass whole genome sequencing (WGS) at ~1x and genotype imputation was performed by Gencove. Low-pass WGS and imputation provides comparable or improved accuracy and variant discovery to array-based genotyping[72,73]. 173 milk samples successfully underwent WGS and imputation from the original 1 mL aliquot DNA extraction. 72 samples had insufficient DNA extracted from the initial 1 mL sample, or failed Gencove’s quality control. Of these 72 samples, 62 had an additional 15 mL frozen aliquot that was shipped to Gencove and DNA was extracted using a mag Nucleic Acid Purification Kit (Biosearch Technologies), and ~1x low-pass WGS was performed. 11 of these samples failed Gencove’s quality control and 51 samples successfully underwent WGS and imputation, resulting in a total of 224 samples with genotype information. 2 participants contributed 2 milk samples (from 2 separate pregnancies), and we included only one sample per participant in our analyses, leaving 222 unique individuals with genotype information. Sample-level details of extraction and sequencing are in Table S1.

BCFtools[74] was used to combine VCFs into a BCF file for all individuals, filtering for minor allele frequency >1% and maximum missing genotypes of 5%. A genetic relatedness matrix was generated with the PLINK[75] (v1.90b6.10) ‘--make-rel’ command, and pairs of individuals with relatedness coefficient >0.05 were pruned, leaving 206 individuals for genetic analyses. Genetic PCs for these 206 individuals were calculated with PLINK using 67,878 SNPs with minor allele frequency >5% after pruning for linkage disequilibrium (PLINK command ‘--indep-pairwise 500 5 0.5’). Genetic ancestry proportion estimates provided by Gencove showed that individuals included in genetic analyses were mostly of European genetic ancestry, with only 17 of 206 individuals with estimated European ancestry <95% (Fig. S2).

We checked for sample swaps by performing genotype calling from RNA-seq reads aligned to chromosome 2 using ‘bcftools mpileup’, and using ‘bcftools gtcheck’ to compare genotypes from RNA-seq to Gencove variant calls from low-pass WGS[74]. We did not detect any sample swaps: for all samples included in eQTL analysis, the DNA sample with matching sample ID had the lowest average concordance, compared to all DNA samples with a different sample ID (Fig. S3).

#### Comparison to GTEx transcriptomes

We downloaded gene-level counts for GTEx samples from the GTEx portal (dataset GTEx_Analysis_2017-06-05_v8_RNASeQCv1.1.9_gene_reads.gct.gz). We filtered to only female GTEx samples, removed tissues with fewer than 19 remaining samples, and then selected 19 random samples for each tissue. We filtered to genes that were detected in both datasets after filtering genes with count 0 across all GTEx & milk samples, leaving 30,468 genes. We then used the thinCounts function in edgeR to downsample each GTEx and milk sample to 5 million read counts. We took the resulting count matrix as a DESeq2 object and performed variance stabilizing transformation (VST). We then took the VST matrix of only GTEx samples, selected the 1000 most variable genes, and performed principal component analysis in R with the 'prcomp' function. We then projected the milk samples onto the PCA scatter plots by calculating each milk sample's values from the GTEx-only PCA.

To compare TPM values across milk and GTEx samples (Fig. 1B), we downloaded gene-level TPM values from the GTEx portal (GTEx_Analysis_2017-06-05_v8_RNASeQCv1.1.9_gene_tpm.gct.gz). We filtered to include only female GTEx samples and filtered to protein-coding genes (as annotated in EnsDb.Hsapiens.v86) and removed histone genes. We then rescaled the TPM for each GTEx and milk sample to again sum to 1 million and calculated each gene's median TPM across a tissue type.

#### Correlations between milk gene expression and maternal/infant traits

We used edgeR[76] to test for correlations between milk gene expression and maternal/milk traits, including all tested traits and technical covariates. Included traits were: Milk CRP concentration, milk glucose concentration, milk IL-6 protein concentration, milk insulin concentration, milk leptin concentration, milk volume expressed, milk fat %, gestational weight gain, milk lactose %, maternal pre-pregnancy BMI, maternal age, and parity (Table S2). We scaled each trait to a mean of zero and standard deviation of one, except binary traits and parity, for which we use the integer number of previous births. The milk RNA-seq count matrix was filtered to include samples with measurements for all tested traits, leaving 158 samples. The count matrix and metadata were loaded into an edgeR object and the "filterByExpr" was used to remove lowly expressed genes, leaving 12,584 genes. We then used the 'estimateDisp' function on a design matrix regressing gene expression across all 12 traits. This model accounted for potential confounding technical effects, including sample mass RNA and sample RIN, by including them as covariates. We then used 'glmQLFit' to fit a quasi-likelihood negative binomial generalized log-linear model to the count matrix and design model, and 'glmQLFTest' to perform a quasi-likelihood F-test of each gene against each tested trait. We used Benjamin-Hochberg correction of P-values across all 12,584 genes x 12 traits.

We tested for gene ontology enrichment of significant genes (q-value<10%) for each trait using the R package topGO[77], with all tested genes as the background gene list. We used the 'resultFisher' function to run a classic Fisher's exact test for each gene ontology, and used a Benjamini-Hochberg correction[78] for all ontologies (N=14,226) across the 9 traits with significant genes (128,034 tests). We report pathways with q-value<10%, fewer than 500 annotated genes, and an overlap of more than 5 genes with the significantly associated gene list for each trait (Table S4). 6 traits (milk glucose, milk IL6, milk insulin, milk volume expressed, milk lactose, and parity) had enriched ontologies that met these criteria.

#### Examination of *PER2* expression and milk traits

Circadian rhythm genes were defined as those in KEGG pathway 'hsa04710'. To test if the time of day of the milk sample collection study visit explained the relationship between *PER2* expression and expressed milk volume, we transformed the time of the study visit into a quantitative variable with the R package 'lubridate'[79]. *PER2* expression values from a variance-stabilizing transformation of the sample-by-gene count matrix in DESeq2[80] were used, including sample RNA mass and RIN as covariates. Regression models were calculated with 'lm' in R. Study time of day was correlated with

*PER2* expression in a simple linear regression (B=0.07, P=0.04), but not with milk volume expression (B=-0.07, P=0.3) We then ran the following linear models:

(A) milk volume ~ *PER2* expression
(B) milk volume ~ *PER2* expression + time of study visit

The linear models were compared with the 'anova' command in R, with a non-significant P-value suggesting that adding the time of study visit variable did not provide a better fit to the data. The same analysis was performed with normalized milk fat percentage replacing milk volume to assess the relationship between milk fat, *PER2* expression, and time of study visit.

#### Deconvolution of bulk transcriptomes with Bisque

Raw gene counts (MIT_Milk_Study_Raw_counts.txt.gz) and metadata (MIT_milk_study_metadata.csv.gz) were downloaded for the Nyquist et al. study[17] from the Broad Insitute Single Cell Portal (https://singlecell.broadinstitute.org/single_cell/study/SCP1671/cellular-and-transcriptional-diversity-over-the-course-of-human-lactation) on 6/3/2022. Count data was filtered to keep just one sample per participant, requiring samples to have been collected >14 days and <3 months postpartum, leaving 10 samples. The count matrix and associated metadata was then put into a the Bioconductor 'ExpressionSet' object format, combining the two macrophage cell type annotations from Nyquist et al. into one cell type called just "Macrophage" and resulting in 8 cell type annotations. The milk gene-level count data was then loaded into an ExpressionSet object and Cell type deconvolution was run with the R package "BisqueRNA" and the function 'ReferenceBasedDecomposition', with parameters "markers=NULL" and "use.overlap=F". Bisque[30] used 27,889 genes present in both the bulk and single-cell expression sets.

#### Milk eQTL analysis

Gene-level quantifications were filtered for the 206 individuals with RNA-seq and genotype data. Genes were filtered to retain those with ≥ 6 counts and and TPM > 0.1 in at least 20% of samples, leaving 17,351 genes of the original 44,747. TPM quantifications were then rank-normalized with the 'RankNorm' function in R package RNOmni[81], and gene coordinates were added using annotations from R package 'EnsDb.Hsapiens.v86'. Genes without coordinate annotations, mitochondrial, and Y chromosome genes were removed, leaving 16,999 genes used in eQTL analyses.

The APEX toolkit was used for cis-eQTL analysis ( https://corbinq.github.io/apex/doc/)[82]. First, 50 latent factors from the gene expression matrix were calculated using command 'apex factor' with 10 iterations. cis eQTL analysis was run with the command 'apex cis' with 5 genetic PCs and 25 gene expression latent factors as covariates, using the LMM model with a genetic relatedness matrix calculated as above in PLINK, and with distance to start site weighting for eGene p-values (ACAT-dTSS). SNPs with minor allele frequency >1%, missing genotype information <5%, and within 1Mb of the gene transcription start site were included. The command used was as follows:

apex cis --bcf [genotypes bcf file] --bed [gene expression bed file] --cov [genetic PCs + gene expr. LFs covariate file] --grm [genetic relatedness matrix] --prefix [output file prefix] --long --dtss-weight 0.00001

APEX uses an aggregated Cauchy association test to calculate a gene-level P-value, and can use the distance to TSS weighting to improve discovery power (parameter '--dtss-weight' in the command above). eGene P-values were adjusted for multiple tests using a Benjamini-Hochberg correction[78].

### Colocalization of milk and GTEx eQTLs

eQTL summary statistics for single tissues (*.v8.signif_variant_gene_pairs.txt.gz), and gene eQTL summary (*.v8.egenes.txt.gz) were downloaded from the GTEx portal (https://gtexportal.org/). Tissues with eQTL sample size less than 100 were excluded. For each gene with an eQTL in milk at q-value<5%, each GTEx tissue with a significant eQTL (q-value<5%) was identified, and colocalization between the milk and GTEx tissue performed with the *coloc* R package[83]. cis-eQTL summary statistics for each tissue were filtered for those present in both milk and GTEx, statistics were harmonized so the reference/alternative alleles matched, then colocalization was run with the command 'coloc.abf'. A milk eQTL-GTEx eQTL was designated as 'colocalized' if the ratio PP.H4/(PP.H4+PP.H3) > 0.8; as 'not-colocalized' if the ratio PP.H3/(PP.H4+PP.H3) > 0.8; and 'ambiguous' otherwise.

To compare the fraction of milk colocalized eQTLs across GTEx tissues, we needed to take into account that the number of eQTLs for a given tissue is a linear function of the sample size[84], and that eQTLs discovered in a smaller sample size are more likely to be those shared across tissues[85]. We observed a clear linear relationship between the number of eGenes included in our colocalization analysis for each GTEx tissue with the fraction of 'colocalized' eQTLs with milk (Fig. S6), and so we used the residual fraction of eQTLs colocalized in this regression to compare across tissues (i.e. in Fig. 2E). Binomial confidence intervals for this fraction were estimated with a normal approximation.

eQTLs were denoted as milk-specific if either of these criteria were met: (1) there were no GTEx tissues with a significant eQTL for the gene (q-value<5%), or (2) there were no tissues with an eQTL that colocalized with milk, and at least 75% of tested tissues' eQTLs were categorized as not-colocalized. Enrichment analysis of genes with milk-specific eQTLs or tissue-shared eQTLs was performed with the enrichR R package[86] with reference database "GO_Biological_Process_2021". Significantly enriched pathways (those with a reported 'Adjusted.P.value'<0.1) were filtered to retain only pathways whose gene overlap with the tested gene set did not intersect any other enriched pathways' overlapping genes, retaining the pathways with the smallest P-values. This approach yielded 2 enriched pathways for each gene set (Fig. 2D).

### Overlap between milk eGenes and dairy cattle QTL

Cattle gene coordinates for ARS_UCD1.2 genome were downloaded from https://bovinegenome.elsiklab.missouri.edu/downloads/ARS-UCD1.2, filtered for mRNAs, and for each gene with multiple entries the entry with the largest region was retained. Dairy cattle QTL were downloaded from the animalQTLdb (https://www.animalgenome.org/cgi-bin/QTLdb/index) by selecting "All data by bp (on ARS_UCD1.2 in bed format)".

For each of 4 milk-related traits, we selected QTL with the following trait labels: milk yield (Milk yield, 305-day milk yield, Average daily milk yield), milk somatic cell count (Somatic cell score, Somatic cell count), milk protein (Milk protein percentage, Milk protein yield, Milk protein content), and milk fat (Milk fat percentage, Milk fat yield, Milk fat content). To identify a smaller list of genes identified in QTL from multiple studies, as some of these traits' QTL overlapped thousands of genes, we identified genes that overlapped at least 1 QTL for all 4 dairy cattle milk traits (N=1,035 genes, Table S7).

To test for enrichment of milk-specific vs. tissue-shared eQTL genes in these lists, we filtered genes for those that were called as milk-specific vs. shared and that were present in the dairy cattle genome annotation above (N=837 genes). We performed a two-sided Fisher's exact test where the 2x2 contingency table axes were: (A) milk-specific vs. tissue-shared eGenes (from our human milk eQTL analysis), and (B) cattle QTL overlapping genes vs. cattle QTL nonoverlapping (from the gene lists identified above), using the 'fisher.test' command in R.

#### Colocalization of milk eQTLs with breast cancer GWAS summary statistics

GWAS summary statistics from Zhang et al. 2020[40] (icogs_onco_gwas_meta_overall_breast_cancer_summary_level_statistics.txt.gz) were downloaded from the BCAC website (https://bcac.ccge.medschl.cam.ac.uk/bcacdata/oncoarray/oncoarray-and-combined-summary-result/gwas-summary-associations-breast-cancer-risk-2020/). Coordinates were converted to hg38 with LiftOver, and the meta-analysis summary statistics for all breast cancers were used (column names 'Beta.Meta', 'p.meta', etc.). For each milk eGene, colocalization was performed if there was a breast cancer GWAS hit of $P < 5x10^{-8}$ within the eQTL window (within 1 Mb of gene TSS). Breast cancer GWAS and milk eQTL summary statistics were filtered to variants within 500 kb of the smallest milk eQTL P-value, and statistics harmonized so the reference/alternative alleles matched, then colocalization was run with the command 'coloc.abf' using the *coloc* R package[83].

#### Correlations between milk gene expression and oligosaccharides

HMOs were rank normalized within the 48 individuals with both gene expression and HMO data, using the 'RankNorm' function from R package 'RNOmni'. For HMOs absent in non-secretors (2'FL and DFLac; Fig. S7), we included only secretor individuals (N=32). The milk RNA-seq count matrix was filtered to include samples with HMO measurements (N=48 for all samples, N=32 for secretors only). The following HMO categories were also calculated: the sum of all HMO concentrations, the sum of all sialylated HMO concentrations (DSLNH, DSLNT, FDSLNH, LSTb, LSTc, 3'SL, 6'SL), and the sum of all fucosylated HMO concentrations (DFLNH, DFLNT, FDSLNH, FLNH, LNFP-I, LNFP-II, LNFP-III, 3'FL, DFLac, 2'FL). These HMO category sums were rank normalized across all individuals.

We used edgeR to test for correlations between milk gene expression and HMO concentrations. The count matrix and metadata were loaded into an edgeR object and "filterByExpr" was used to remove lowly expressed genes, leaving 12,471 genes. For each HMO, we then used the 'estimateDisp' function on a design matrix regressing gene expression across HMO concentration, sample mass RNA, and sample RIN. We then used 'glmQLFit' to fit a quasi-likelihood negative binomial generalized log-linear model to the count matrix and design model, and 'glmQLFTest' to perform a quasi-likelihood F-test of each gene against each tested HMO. We used Benjamin-Hochberg[78] correction of P-values across all HMO-gene pairs.

We tested for gene ontology enrichment of significant genes (q-value<10%) for each trait using the R package topGO, with all tested genes as the background gene list. We used the 'resultFisher' function to run a Fisher's exact test for each gene ontology, and used a Benjamini-Hochberg correction[78] for all ontologies (N=14,178) across the 6 HMOs/HMO categories with at least 10 significant genes (DSLNT, LSTb, LSTc, 3'SL, sialylated sum, all HMOs sum). We report pathways enriched with q-value<10% for each HMO/HMO category (Table S11).

#### Genetic associations at milk eQTLs with milk oligosaccharides

The list of candidate genes to test for effects of milk eQTLs on HMO concentrations was downloaded from Supplementary Dataset 2 in Kellman et al, 2022[42]. From this gene list, we identified 8 genes with significant eQTLs in our dataset (q-value<10%). To test for genetic associations between the lead SNP (smallest P-value) at milk eQTLs and HMO concentrations using the rank-normalized HMO concentrations described above. For 2'FL and DFLac, which were absent in non-secretors (Fig. S7), we rank-normalized the concentrations within secretors and scaled concentrations in non-secretors to have mean -3 and s.d. 0.1, to avoid introducing variation that did not exist in non-secretors. We used 'glm' in R to fit a model with HMO concentrations as the outcome, including genotype, secretor status and the first five genetic PCs as covariates:

HMO ~ genotype + secretorStatus + PC1 + PC2 + PC3 + PC4 + PC5

For models of HMOs vs. *FUT2* eQTL genotype, we excluded the secretor status term.

To test for potential causal effects between milk gene expression and HMO concentrations, we used a Wald Ratio, which estimates the causal effect between an exposure (milk gene expression) and outcome (HMO concentration) by dividing a single genetic variant's effect on outcome by the genetic effect on the exposure[87]. We used the Wald Ratio rather than a two-stage least squares approach because we had a much larger sample size for our eQTL analysis (N=206) than HMOs (N=41), and we wanted to leverage the eQTL effect estimates from this larger sample size. Two-stage least squares, which is used in a one-sample study design such as ours, would require the same participants to have both exposure and outcome data, limiting the sample size for estimates of genetic variant effects on gene expression to N=41.

#### Processing of infant fecal metagenomes

Infant fecal collection and storage, and metagenomic DNA extraction were described previously[63]. Briefly, feces were collected from diapers either during study visits and frozen at -80°C immediately, or collected at home, stored in 2 ml cryovials with 600 µl RNALater (Ambion/Invitrogen, Carlsbad, CA), and stored at -80°C after shipping to the lab at the University of Minnesota. DNA was extracted using the PowerSoil kit (QIAGEN, Germantown, MD), eluted with 100 µl of the provided elution solution, and stored in microfuge tubes at -80°C.

Extracted DNA was used to construct libraries for metagenomics sequencing using the Illumina Nextera XT ¼ kit (Illumina, San Diego, CA, United States). Metagenomics libraries were then sequenced on an

Illumina NovaSeq system (Illumina, San Diego, CA) using the S4 flow cell with the 2x150 bp paired end V4 chemistry kit by the University of Minnesota Genomics Center, achieving a sequencing depth of ~4.5 million reads per sample.

Microbial taxon abundances were generated by first processing metagenomic fastq files with Shi7 version 1.0.1 (Al-Ghalith, et al., 2018), which learns optimal quality control parameters from the data. Sequences were then trimmed, filtered by quality scores, and stitched per the learned parameters in Shi7. Sequences from all samples were multiplexed into a single fasta file for downstream processing. Processed sequences were aligned to reference databases using BURST version 0.99.7f, (Al-Ghalith and Knights, 2017) using a reference genome database generated from GTDB r95 (https://gtdb.ecogenomic.org/stats/r95). A 95% identity cutoff and forward/reverse complement flag were used. Resulting .b6 files were converted to reference and taxonomy tables using embalmulate (Al-Ghalith and Knights, 2017) with 'GGtrim' activated. To generate microbial pathway abundances, metagenomic sequences were run through the MetaPhlAn version 3.0.7 pipeline, with BowTie2 version 2.4.2 64-bit, DIAMOND version 0.9.24, and MinPath version 1.5.

To generate the PCA of infant metagenomes in Fig. 4A, data were filtered to include only taxa with relative abundance >0.001 in at least 10% of 1-month or 6-month samples. A centered log-ratio transformation was performed on the relative abundances of each sample, and principal components were calculated with the 'prcomp' command in R.

#### Sparse CCA of human milk transcriptomes and infant fecal metagenomes

Input datasets were prepared as follows:

**Milk gene expression**: To prepare gene expression data for this analysis, the sample-by-gene count matrix was loaded into DESeq2[80], filtered to keep only protein-coding genes with count>0 in at least half the participants (13,157 genes), and transformed using the variance stabilizing transformation. After this transformation, the variance of each gene was calculated across all samples and genes in the lowest 25% variance were removed, leaving 9,868 genes.

**Infant fecal metagenomes**: Taxon abundances and pathway abundances from 1-month infant fecal samples were processed separately. The taxon relative abundance matrix was filtered to retain species-level taxa only, keeping only 96 species with a relative abundance $>1\times10^{-3}$ in at least 10% of samples. A centered log-ratio transformation was then performed on each sample's relative abundances. For microbial pathways, species-specific and unclassified pathways were removed, leaving 145 pathways. The species and pathway level information was then combined into one matrix.

Each dataset was filtered for the 108 individuals with both 1-month infant fecal metagenomes and 1-month milk gene expression. Sparse canonical correlation analysis (sparse CCA), to identify sparse components maximizing correlation between the milk gene expression and infant fecal metagenome datasets, and enrichment analyses of genes in each sparse component, were performed as previously described[88], using k=15 components. Code was downloaded from https://github.com/blekhmanlab/host_gene_microbiome_interactions. Significance of the sparse components was calculated with leave-one-out cross-validation, and 10 components were retained at

Benjamini-Hochberg q-value<10%. Pruning significant components whose scores across mother-infant pairs were correlated at Pearson's r>0.75 left 6 remaining sparse components (Fig. 4B).

To generate network interaction plots between milk-expressed genes and infant fecal microbes identified in the sparse CCA analysis, for each significantly enriched pathway (q-value<10%) in a component, we (1) filtered for overlapping genes between the component and pathway; (2) generated a pairwise correlation matrix of mother-infant pairs' trait values for those genes, the top 3 microbiome traits in the component with positive weights, and top 3 microbiome traits with negative weights; (3) pruned for correlations with Pearson's r>0.3; (4) generated a network plot from the pairwise correlation matrix using the 'ggnetwork' package in R[89].

*B. infantis* growth rates were estimated using Compute PTR (CoPTR)[90]. We aligned the infant gut metagenomic shotgun reads to the *B. infantis* ATCC 15697 reference genome, downloaded from NCBI, using bowtie2 v2.2.4[91]. We then used CoPTR to get coverage information for each mapped sample, filtering for samples with at least 75% coverage and at least 3000 mapped reads to the *B. infantis* genome. For samples that passed these filters, CoPTR was used to estimate the peak-to-trough ratio (PTR) from the coverage information, an estimate of the bacterial growth rate.

#### Associations between maternal genotype at eQTLs and HMOs or the infant fecal microbiome

For each milk-specific eQTL, a tag SNP was defined as that with the smallest P-value, requiring there to be at least 10 individuals in the smallest diploid genotype class. 1- and 6-month infant microbiome taxa abundances and pathway abundances were processed as described above for sparse CCA. Microbiome data was retained for mother-infant pairs with maternal genotype data and 1- or 6-month metagenomic data, leaving N=105 mother-infant pairs. Within each time point and microbial taxon or pathway dataset, microbiome traits were pruned by removing one trait from trait pairs with the highest Pearson correlation until no pairs remained with r>0.75. As we were hoping to detect genetic effects from gene expression in milk, we regressed microbiome traits against infant delivery mode (cesarean vs. vaginal), and for 6-month traits whether the mother-infant pair was exclusively breastfeeding, and if complementary foods were introduced. At 1 month postpartum all pairs were exclusively breastfeeding with no complementary foods. We also regressed microbiome traits against maternal 10 genetic PCs, using the residuals of the following regression models using 'glm' in R:

1-month microbiome trait ~ delivery_mode[0/1] + PC1 + PC2 + … + PC10

6-month microbiome trait ~ delivery_mode[0/1] + exclusive_breastfeeding[0/1] +
complementary_foods[0/1] + PC1 + PC2 + … + PC10

We then performed genetic association testing of maternal genotype at each milk-specific eQTL tag SNP against each infant microbiome trait, using a linear mixed model Wald test in GEMMA v0.98.5[92]. There were 232 unique eQTL tag SNPs tested against 313 microbiome traits (46 1-month taxa, 103 1-month pathways, 58 6-month taxa, 104 6-month pathways) (Table S16).

### Supplemental figures

#### Figure S1

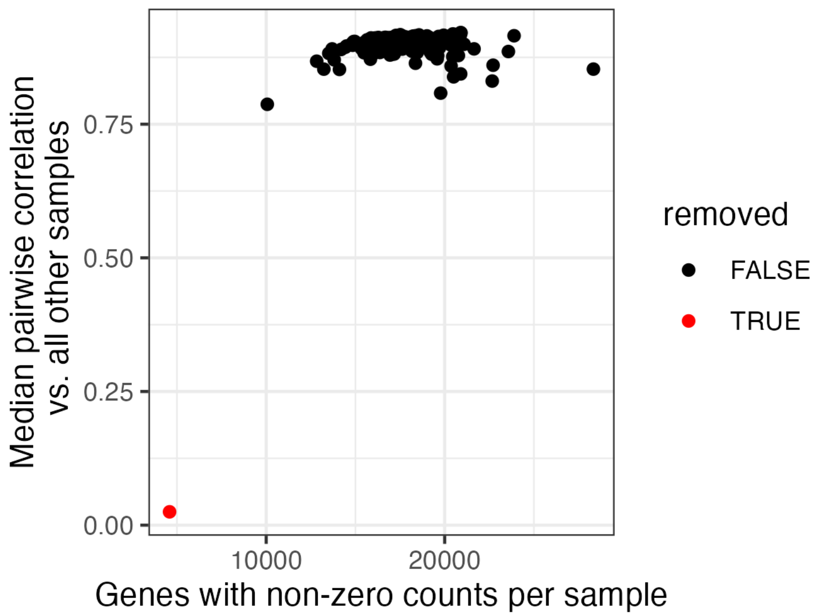


Distribution of median pairwise correlations of TPM counts for each sample against all other samples (y-axis), and the number of genes with non-zero counts per sample in RNA-seq data. The red dot indicates an outlier sample that was removed from further analysis.

Figure S2

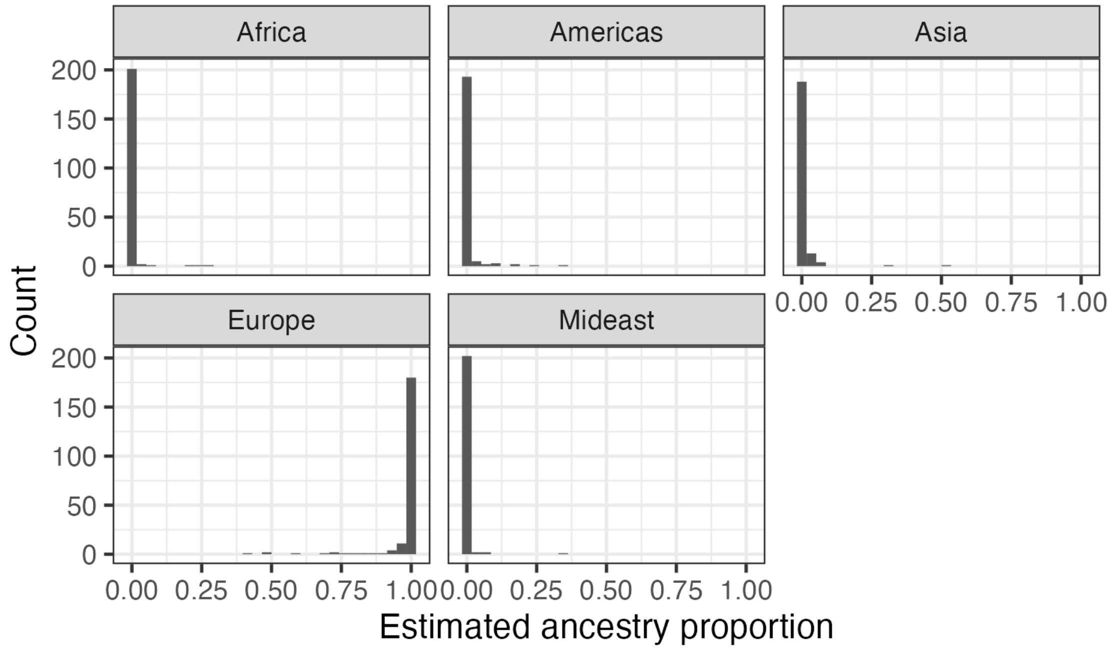


Distributions of genetic ancestry estimates for individuals included in the eQTL analysis. Within each panel, representing a continental ancestry group, is a histogram displaying the distribution of estimated ancestry proportions for that group for all samples. e.g. all samples have an estimated European ancestry proportion >0.4, with the majority ~1; while no samples have estimated African ancestry proportion > 0.3.

Figure S3

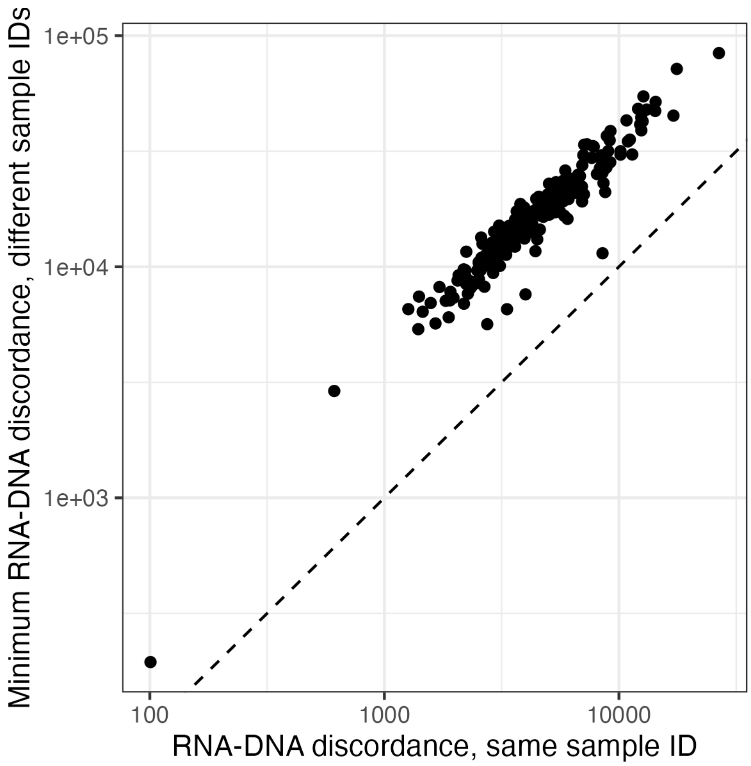


Distribution of discordance between genotypes estimated from RNA and DNA samples. Each dot represents a single sample ID, with the x-axis showing the genotype discord between genotype calls between the RNA and DNA samples with the same sample ID. The y-axis is the minimum discord between that sample ID's RNA sample and any DNA sample. All points are above the x=y line (dashed line), showing that the DNA sample with the matching sample ID always had the most similar genotype calls for each RNA sample, and that there were no sample label mix-ups.

Figure S4

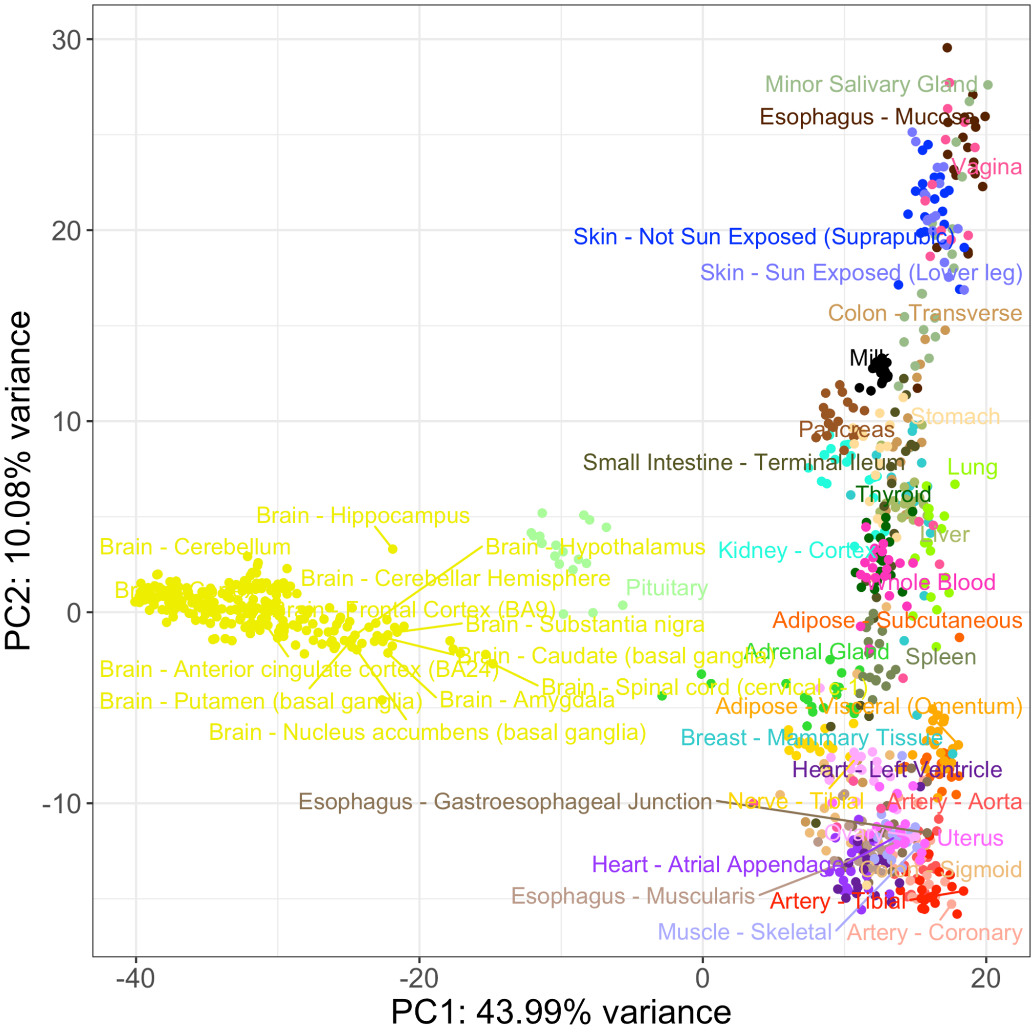


Principal component analysis of transcriptomes from a subset of GTEx tissues and milk. PCs were calculated using the 1000 most variable genes within GTEx, then milk samples were projected onto the GTEx samples.

#### Figure S5

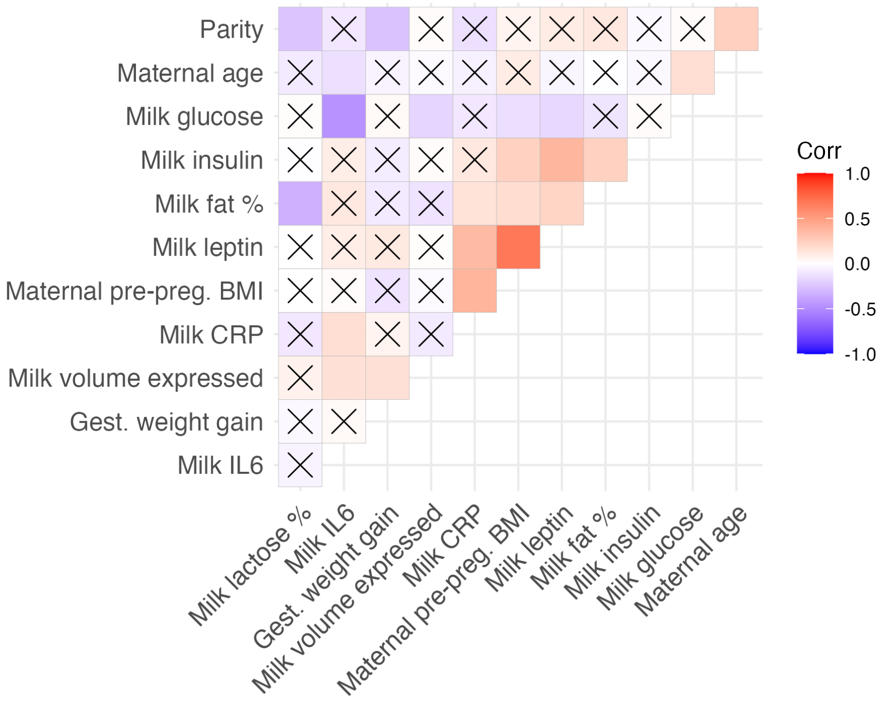


Spearman correlations among the 12 maternal/milk traits tested for relationships with milk gene expression. An "X" signifies P>0.05, i.e. not significant.

Figure S6

(A)

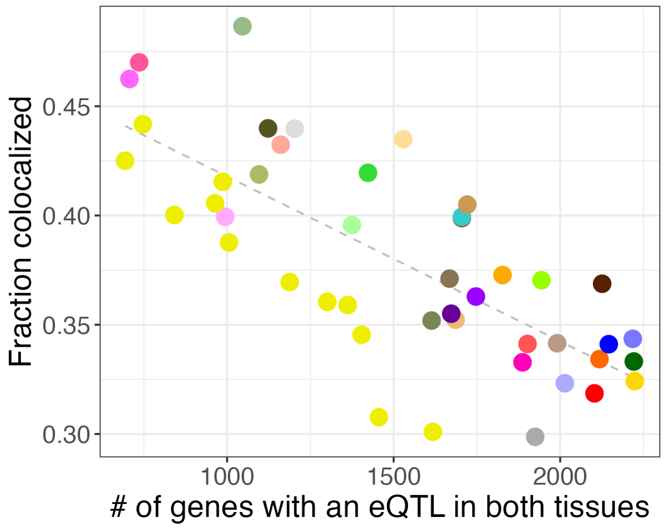


(B)

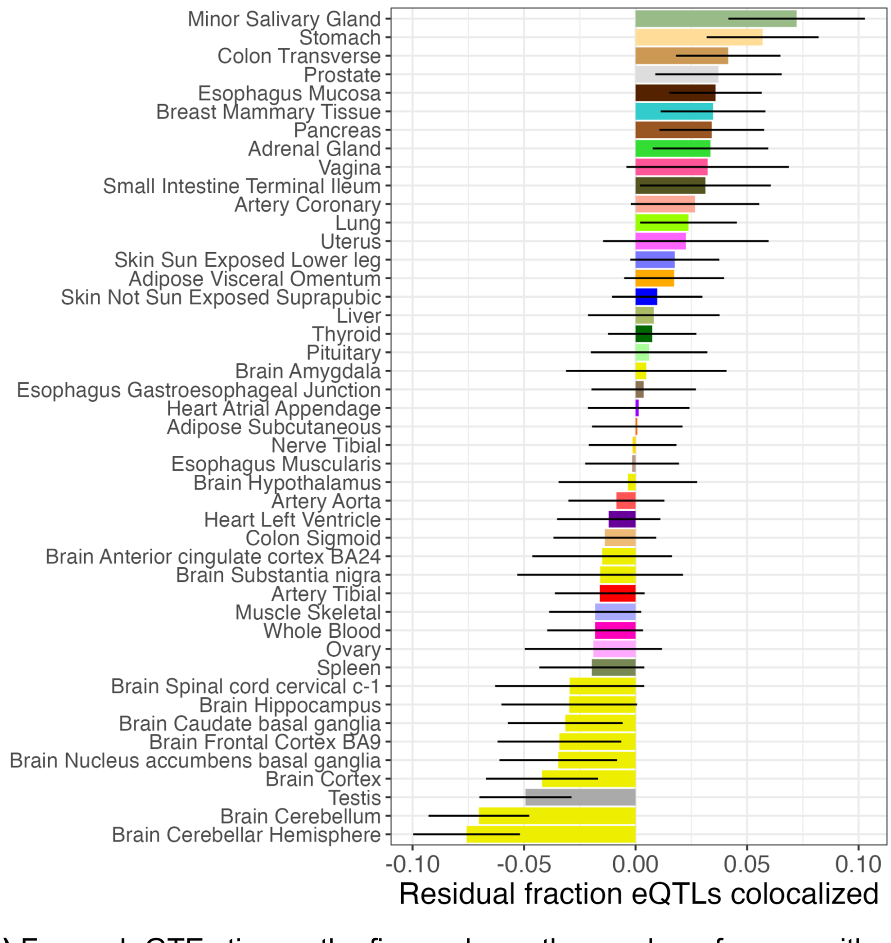


**(A)** For each GTEx tissue, the figure shows the number of genes with eQTLs in the tissue and milk (x-axis), vs. the fraction of colocalized eQTLs between milk and the given tissue (y-axis). The dashed line illustrates a significant linear regression slope ($P = 2.7x10^{-9}$). **(B)** Sharing of eQTLs between milk and 45 GTEx tissues, measured through statistical colocalization. Each bar shows each tissue's similarity to milk, measured by residual fraction of eQTLs colocalized with milk, after regressing out tissue sample size. Error bars represent a 95% confidence interval.

#### Figure S7

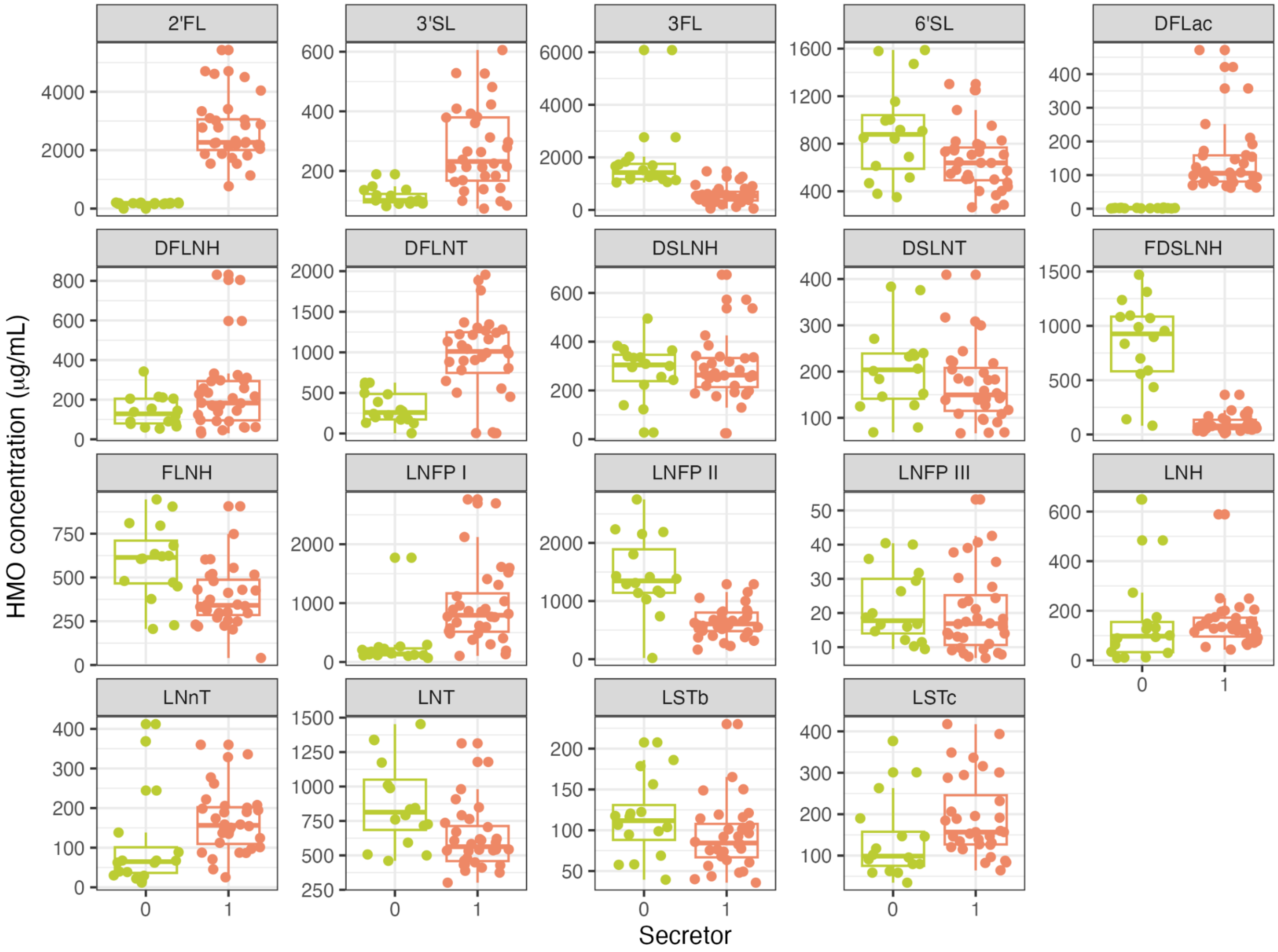


Distributions of HMO concentrations, grouped by secretor (orange, N=32) and non-secretor (light green, N=16) individuals.

Figure S8

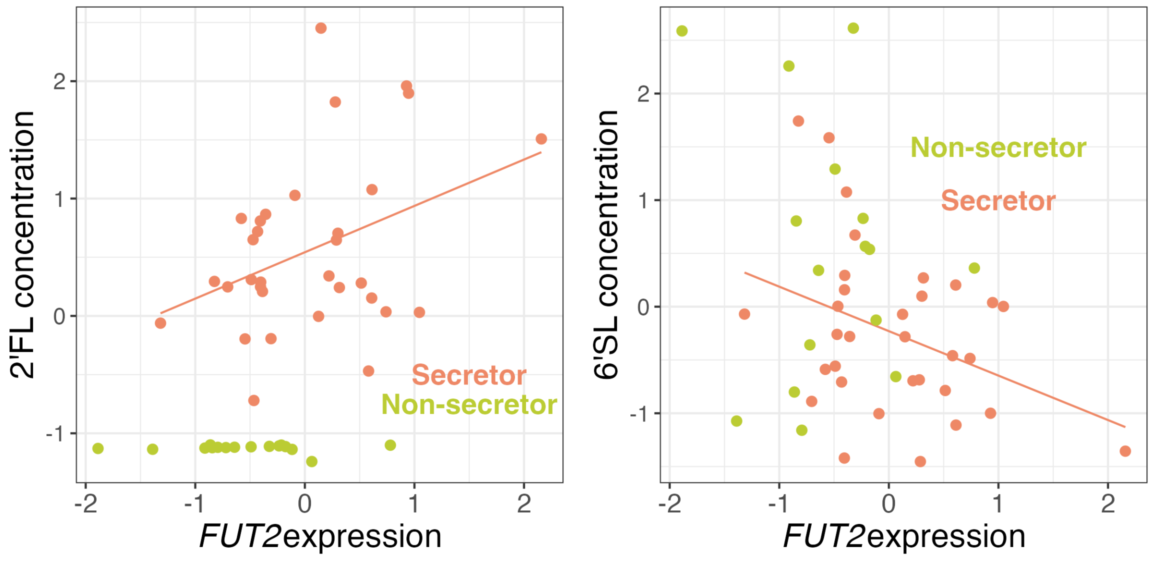


Associations between normalized *FUT2* expression (x-axis) and the normalized concentration (y-axis) of two HMOs (2’FL: Beta = 0.40, P = 0.03; 6’SL: Beta = -0.42, P = 0.04). Regression statistics are for secretor individuals only (N=32). Secretors are shown in orange and non-secretors in light green.

### Figure S9

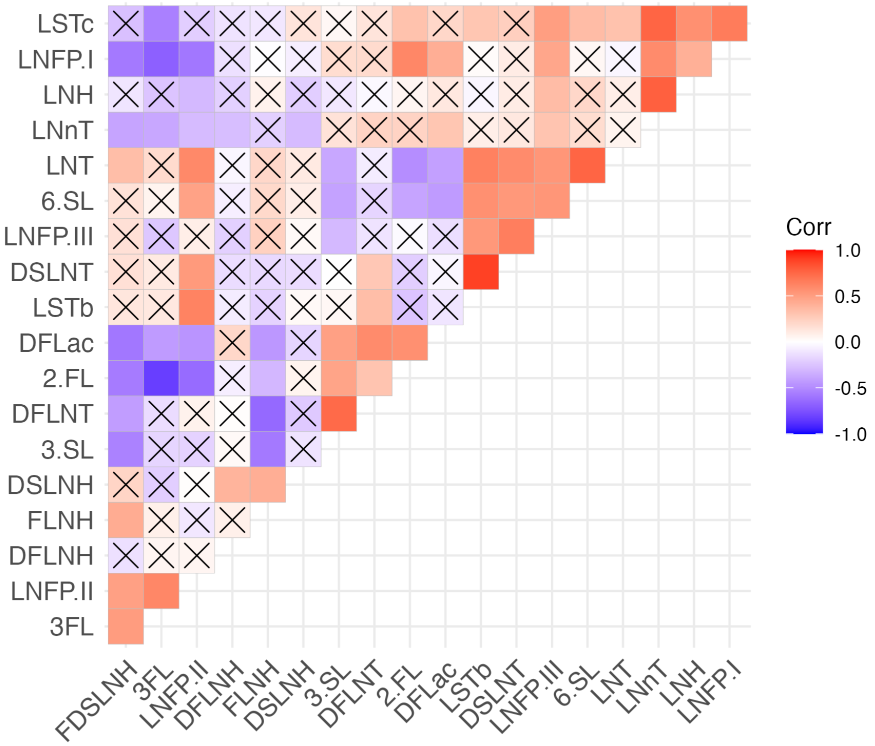


Pairwise Pearson correlations between rank-normal-transformed HMO concentrations. An “X” signifies q-value>0.1, i.e. not significant.

Figure S10

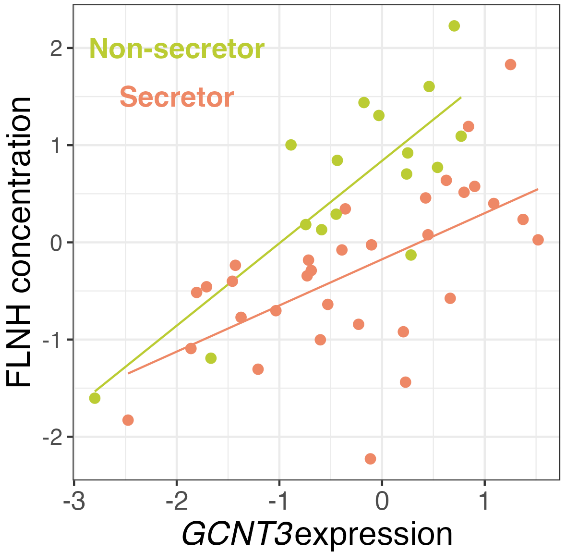


Correlation between normalized *GCTN3* expression and normalized FLNH concentration. Secretors are shown in orange and non-secretors in light green. To visualize the positive correlation in both secretors and non-secretors, regression lines are shown for secretors and non-secretors separately, but the relationship was assessed using all individuals with secretor status as a covariate in edgeR as described in Materials and Methods. Beta= 0.67, P=1.26x10$^{-6}$, q-value=0.04.

Figure S11

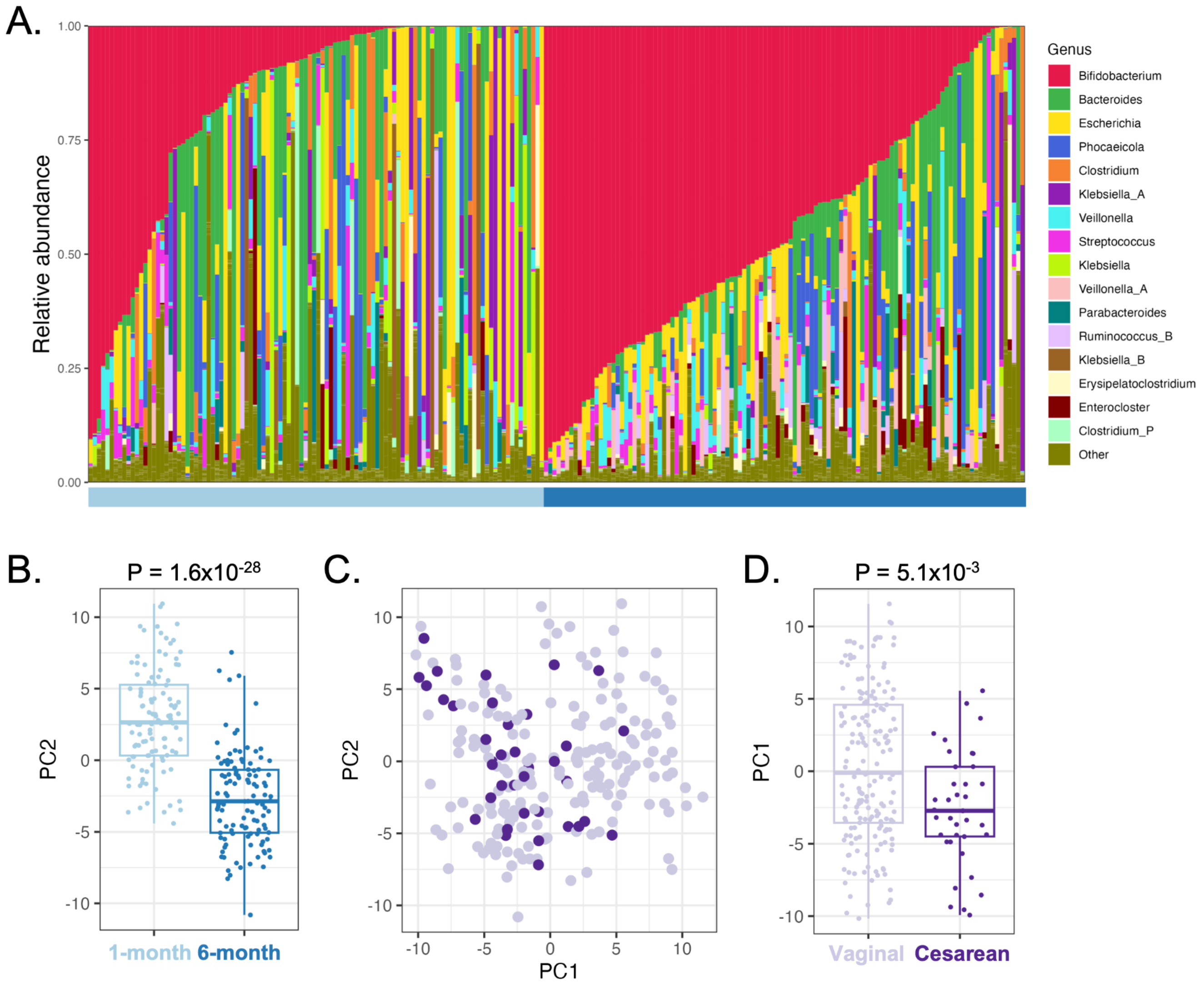


Infant metagenomic data summarized at the taxonomic level. **(A)** Barplots showing the relative abundances of bacterial genera, grouped by 1-month (light blue, N=108) and 6-month (dark blue, N=113) samples. Each bar represents a sample. **(B)** Values of PC2 (y-axis; principal component 2 in Fig 4A) grouped by sample time point (x-axis). There was a significant difference between the two timepoints, using a linear mixed effects model with sample time point and delivery mode as fixed effects and subject ID as a random effect (timepoint: est. = -5.5, P = $1.6x10^{-28}$). **(C)** Scatter plot of PC1 vs. PC2, as in Fig. 4A, but colored by delivery mode: vaginal (light purple) or cesarean (dark purple). Both 1-month and 6-month samples are plotted. **(D)** Values of PC1 (y-axis; principal component 1 in Figs. 4A and S6C) grouped by delivery mode (x-axis). There was a significant difference in PC1 score between the two delivery mode groups, using a linear mixed effects model with sample time point and delivery mode as fixed effects and subject ID as a random effect (delivery mode: est. = -3.2, P = $5.1x10^{-3}$).

### Supplemental table descriptions

#### Table S1

Summary of samples used in each analysis, and sequencing metrics. study_id: unique participant ID; DNA_conc: concentration of DNA in milk sample (ng/ul). RNA_conc: concentration of RNA in milk sample (ng/ul). RIN: RIN score; expr.inc: logical statement, was sample included in gene expression analyses; eqtl.incl: logical statement, was sample included in eQTL analyses; redo.DNA: logical statement, was DNA extraction & sequencing performed on a larger 10 mL milk aliquot; gt.incl: logical statement, was sample successfully genotyped; inf.fecal.1mo: logical statement, was sample included in analysis with 1-month infant fecal metagenome; inf.fecal.6mo: logical statement, was sample included in analysis with 6-month infant fecal metagenome; WGS_effective_coverage_min: Gencove effective coverage; WGS_fraction_contamination_max: Gencove estimated sample contamination; WGS_snps_min: Gencove number of called SNPs; RNAreadCount: number of RNA-seq reads; RNAuniqMapPct: percentage of uniquely mapping RNA reads, output from STAR; Notes: only present noting which 4 samples come from the 2 participants that contributed two milk samples.

#### Table S2

Overview of MILK study traits used in DEG analysis. trait: trait; N: sample size of trait for normalization (for DEG analysis, only samples with all trait info were used, N=158); Mean: sample mean, Median: sample median, Min: sample minimum; Max: sample maximum; pct2.5: sample 2.5 percentile; pct97.5: sample 97.5 percentile; units: units of measurement.

#### Table S3

Output of correlation analysis between maternal/milk traits and gene expression in milk in EdgeR. Only nominally significant ($P<0.05$) associations are reported. logFC: log2 fold change in gene expression per unit 1 change in trait value; logCPM: log2 counts per million; F: quasi-likelihood F-test statistic; PValue: P-value; trait: trait tested; gene_id; Ensembl gene ID of gene tested; gene_name: gene name of gene tested; all.FDR: Benjamini-Hochberg q-value across all tested trait-gene pairs.

#### Table S4

Output of enrichment analysis in TopGO for significant genes in correlation analysis between maternal/milk traits and milk gene expression. GO.ID: gene ontology ID; Term: gene ontology name; Annotated: number of annotated GO genes included in background gene list; Significant: number of GO genes significantly correlated with the tested trait (q-value<10%); Expected: expected number of significant genes in GO gene list; trait: tested trait; p: Fisher' test P-value; FDR.all: Benjamini-Hochberg q-value corrected across all tested GO terms and traits.

#### Table S5

Spearman correlations between 12 maternal/milk traits and 8 Bisque-estimated cell type proportions. cellType: cell type; trait: maternal/milk trait; r: Spearman correlation coefficient; P: Spearman correlation coefficient P-value; FDR: Benjamini-Hochberg q-value across all cell type/trait pairs.

#### Table S6

Summary of eQTL analysis across 16,999 tested genes. Gene: Ensembl gene ID; gene_name: gene name; gene_biotype: gene biotype; n_samples: number of samples in eQTL analysis; n_cis_variants: number of nearby (cis) genetic variants tested for association with gene expression; egene_pval: gene-level aggregated Cauchy association test (ACAT) P-value (see Methods); FDR: Benjamini-Hochberg q-value across all tested genes; milkSpecific: logical, was eQTL identified as milk-specific in GTEx eQTL colocalization analysis.

#### Table S7

List of genes overlapping QTL for all four dairy cattle milk traits. Used to test for enrichment of dairy cattle milk trait QTL genes in milk-specific vs. tissue-shared eQTLs.

#### Table S8

Breast cancer GWAS loci tested for colocalization with milk eQTL. nsnps: number of SNPs in both milk eQTL & GWAS summary statistics used in coloc analysis; PP.H3.abf: coloc posterior probability of hypothesis 3 (both traits have causal variant, not a shared causal variant); PP.H4.abf: coloc posterior probability of hypothesis 4 (both traits have a shared causal variant); gene: Ensembl gene ID; gene_name: gene name; milk_specific: logical, was milk eQTL identified as milk-specific in colocalization analysis with GTEx eQTLs; Beesley/Ferreira/Fachal/Zhang: logical, was this gene connected to a GWAS locus in the cited functional analyses.

#### Table S9

HMO abbreviations and full names.

#### Table S10

Output of correlation analysis between HMO concentrations and gene expression in milk in EdgeR. Only nominally significant ($P<0.05$) associations are reported. logFC: log2 fold change in gene expression per unit 1 change in HMO concentration; logCPM: log2 counts per million; F: quasi-likelihood F-test statistic; PValue: P-value; HMO: HMO or HMO category tested; gene_id; Ensembl gene ID of gene tested; gene_name: gene name of gene tested; FDR_all: Benjamini-Hochberg q-value across all tested HMO-gene pairs.

#### Table S11

Output of enrichment analysis in TopGO for significant genes in correlation analysis between HMOs and milk gene expression. GO.ID: gene ontology ID; Term: gene ontology name; Annotated: number of annotated GO genes included in background gene list; Significant: number of GO genes significantly correlated with the tested trait (q-value<10%); Expected: expected number of significant genes in GO gene list; HMO: tested HMO/HMO category; p: Fisher' test P-value; FDR.all: Benjamini-Hochberg q-value corrected across all tested GO terms and HMOs/HMO categories.

#### Table S12

Starting from a list of 54 candidate glycosyltransferase genes[42], eight genes had for 8 of these genes with significant milk eQTLs in our data. gene_name: gene name; gene: Ensembl gene ID; egene_pval: eQTL gene-level P-value if gene was included in eQTL analysis; FDR: eQTL gene-level q-value if gene was included in eQTL analysis; median_tpm: median transcript per million of gene in our milk transcriptomes, NA if not included in eQTL analysis; chrom: chromosome of eQTL tag SNP; pos: base position of eQTL tag SNP; ID: SNP ID of eQTL tag SNP; ref: reference allele of eQTL tag SNP; alt: alternative allele of eQTL tag SNP; beta: estimated SNP effect on gene expression; se: standard error of SNP effect on gene expression; pval: P-value of SNP effect on gene expression.

#### Table S13

Genetic associations between candidate HMO gene eQTL tag genetic variations and HMO concentrations, and Wald Ratio estimates of the effect of genetically modified gene expression on HMO concentration. gene_name: gene name; HMO: tested HMO; ga.est: estimated SNP effect on HMO concentration; ga.se: standard error of SNP effect on HMO concentration; ga.p: P-value of SNP effect on HMO concentration; ga.FDR: Benjamini-Hochberg q-value of SNP effect on HMO concentration; wr.b: Wald ratio estimate of the effect of genetically modified gene expression on HMO concentration; wr.se: standard error of Wald ratio effect estimate; wr.p: P-value of Wald ratio effect estimate; wr.FDR: Benjamini-Hochberg q-value corrected Wald ratio P-value.

#### Table S14

Sparse CCA output weights for identified sparse components containing milk-expressed genes and infant fecal microbial taxa or pathways. feature: feature name; weight: weight of feature in sparse component; type: is feature a milk gene (gene) or infant fecal microbial trait (microbe); component: the sparse component this feature weight is for.

#### Table S15

Pairwise Pearson correlations between expression levels of JAK/STAT pathway genes in milk, infant fecal *B. infantis* growth rate and relative abundance, and milk IL-6 concentration. t1: trait 1; t2: trait 2; r: correlation coefficient; P: correlation P-value; N: sample size of correlation estimate; FDR: Benjamini-Hochberg q-value.

#### Table S16

Results of genetic association tested between maternal genotype at milk-specific eQTL tag SNPs and infant fecal microbial taxa or pathways. pt: infant fecal microbiome phenotype; rs: eQTL tag SNP ID, chr: eQTL tag SNP chromosome; ps: eQTL tag SNP base position (hg38); allele1: alternative allele; allele0: reference allele; af: sample allele frequency; beta: estimated SNP effect on phenotype; se: standard error of SNP effect estimate; p_wald: Wald test P-value; FDR: Benjamini-Hochberg q-value across all SNP/phenotype pairs; gene: eQTL Ensembl gene ID; gene_name: eQTL gene name.
